# Symbiotic diversification relies on an ancestral gene network in plants

**DOI:** 10.1101/2025.04.01.646537

**Authors:** Leonardo Castanedo, Katharina Melkonian, Tatiana Vernié, Lucie Chauderon, Tifenn Pellen, Matheus E. Bianconi, Corinne Lefort, Cyril Libourel, Aurélie Le Ru, Simon Aziz, Cristina Latorre, Océane Thiercelin, Nathalie Rodde, Stephane Cauet, Caroline Callot, Laura Forrest, Tomoaki Nishiyama, Wickell D David, Fay-Wei Li, Jean Keller, Pierre-Marc Delaux

## Abstract

Symbioses have been fundamental to colonization of terrestrial ecosystems by plants and their evolution. Emergence of the ancient arbuscular mycorrhizal symbiosis was followed by the diversification of alternative intracellular symbioses, such as the ericoid mycorrhizae (ErM). We aimed at understanding how these diversifications occurred. We sequenced the genomes of ErM-forming liverworts, and reconstituted symbiosis under laboratory conditions. We demonstrated the existence of a nutrient-regulated symbiotic state that enables ErM and underlies intracellular colonization of plant tissues. Comparative transcriptomic analyses identified an ancestral gene module associated with intracellular symbiosis beyond ErM. Genetic manipulations in the liverwort *Marchantia paleacea*, phylogenetics and transactivation assays demonstrated its essential function for intracellular symbiosis. We conclude that plant have maintained, and convergently recruited, an ancestral gene module for intracellular symbioses.

## Main Text

Symbiotic interactions have played a fundamental role in the colonization of terrestrial ecosystems by plants^1–3^, influencing both population dynamics and biogeographical distribution, and shaping plant evolution^3^. The appearance of the first intracellularly-hosted fungal symbiont, the arbuscular mycorrhizal symbiosis (AMS), at the dawn of the land plants 450 million years ago is supported by the fossil record, evo-devo and phylogenetic studies^4–9^. The evolution of AMS in land plants was followed by the diversification of alternative intracellular symbioses and the origin of novel plant clades that colonized previously unoccupied terrestrial ecosystems^3^. For example, the evolution of ericoid mycorrhizae (ErM) around 140–75 Ma coincided with the origin and the appearance of the Ericaceae family of vascular plants^3,10,11^. ErM can also be found in bryophytes, within the leafy liverwort order Jungermanniales (Jungermanniopsida)^12,13^. It is estimated that ErM in Jungermanniales, alongside Ericaceae plants, contribute to the mobilization of nitrogen (N) and phosphate (P) in the habitats where these plants often co-exist^14–16^.

The establishment of AMS relies on a set of four genes (*SYMRK, EPP1, CCaMK* and *CYCLOPS*), which constitute the Common Symbiosis Pathway (CSP)^17–19^. The CSP genes are both conserved and required for the establishment of AMS across land plants, including angiosperms and bryophytes^20,4,6,18,19^. Consistent with this, plant lineages that have lost the ability to form AMS also lost the CSP during evolution^4,21^. Very few exceptions to these correlated losses have been observed, and they always correspond to the occurrence of another type of intracellular symbiosis, such as the nitrogen-fixing root nodule symbiosis in *Lupinus*^5^, or ErM in Ericaceae and Jungermanniales^5,19^. Altogether, this suggests that ErM, like other type of intracellular symbioses, have diversified from the ancestral AMS. However, the cellular, molecular and genetic mechanisms underlying these diversification events remain unclear. Here, we aimed at exploring the mechanisms underlying symbiosis diversification by studying ErM in the leafy liverwort *Calypogeia fissa*.

### Nutrient-dependent mutualistic intracellular symbiosis in *Calypogeia fissa*

We aimed at developing a model for studying ErM in controlled conditions. To do this, we selected the leafy liverwort *Calypogeia fissa* and generated axenic cultures. *C. fissa* was then mock-inoculated, or inoculated with the ErM-forming fungus *Hyaloscypha hepaticicola*, in a nutrient regime low in both phosphate (P) and nitrogen (N) (Fig. 1). We used this treatment to mimic the nutrient-limited conditions in which leafy liverworts can thrive^14^. Two parameters indicative of mutualism were scored: i) hyphal coils in colonized rhizoid-tips which are considered the symbiotic structures^12,22^, and ii) plant growth (Fig. 1A-L). The growth of *H. hepaticicola* was discrete and highly localized around and inside swollen rhizoid tips of *C. fissa* where it formed intracellular coils (Fig. 1G-L). To assess whether colonization benefited *C. fissa* plants, we quantified plant growth in both *H. hepaticicola*-inoculated and mock-inoculated conditions over a 9-week time-course experiment (Fig. 1A-F). We observed that inoculated *C. fissa* plants produced significantly more biomass compared to mock-inoculated plants (Fig. 1C,F). Similar observations were made under extreme nutrient depletion (Fig. S1).

**Figure 1.**
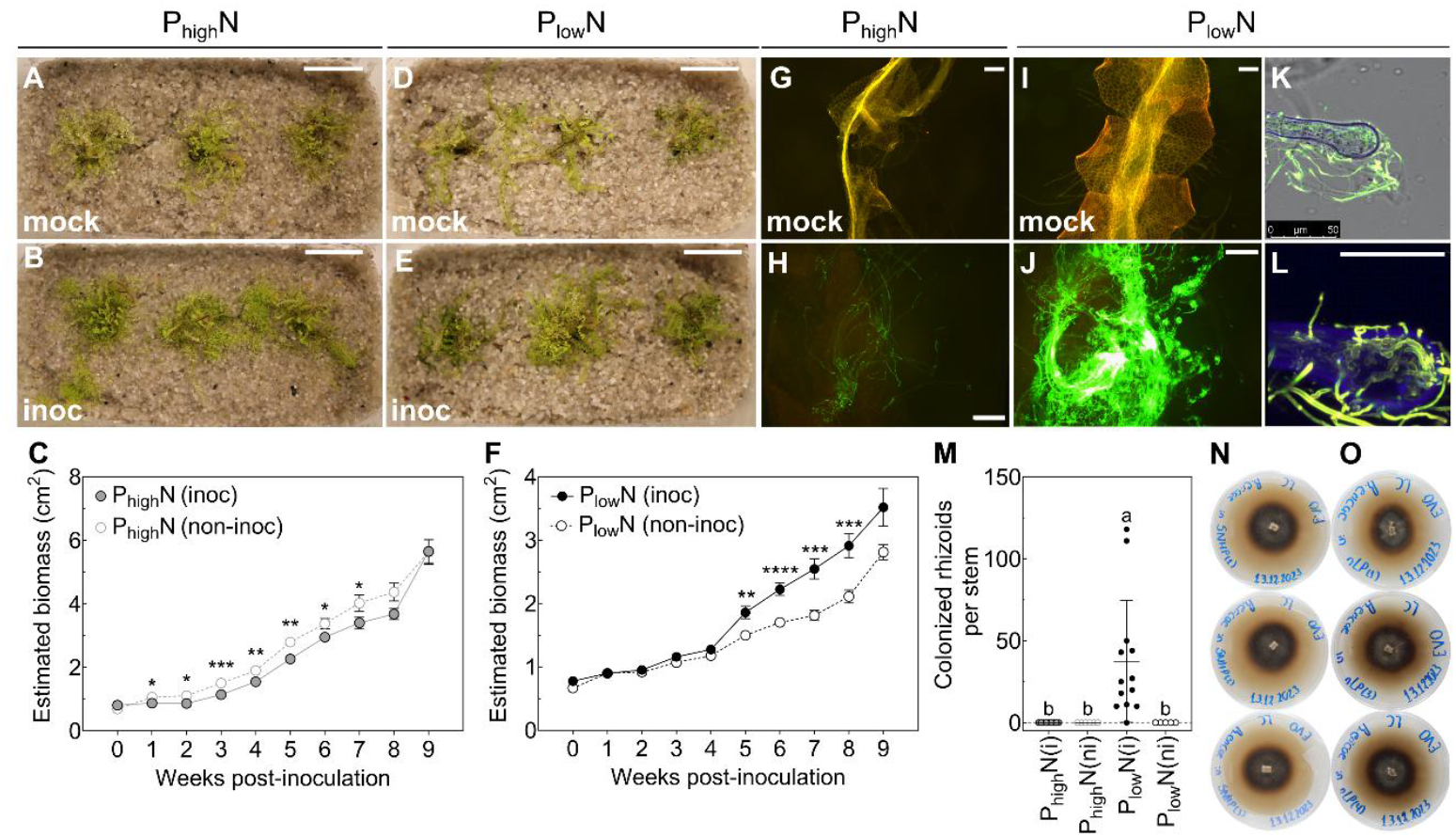
Establishment of Ericoid mycorrhizae in *Calypogeia fissa* relies on a nutrient starvation-induced permissive state. **(A-F)** Representative images (A, B, D, E) and estimated growth (C, F) of *Calypogeia fissa* plants in mock-inoculated (non-inoc) and inoculated (inoc) treatments. Scale bars: 2 cm. Data points in (C, F) indicate the mean ± SEM (*n* = 27) of technical replicates per time-point from one representative experiment of a total of three independent experiments. **(G-L)** Representative fluorescence and confocal microscopy images of *C. fissa* stems harvested from non-inoc (A, D) and inoc (B, E) treatments, and stained with WGA. Plants were photographed and harvested after 8 weeks of growth. Scale bars: 200 μm Confocal microscopy images of samples shown in (K, L), were further stained with Congo red (blue). Scale bar: 25 μm. **(M)** Number of swollen and colonized rhizoid tips per stem in non-inoc (ni) and inoc (i) treatments. Data points indicate the mean ± SD (*n* = 4 to 13) of technical stem replicates from one experiment. Plants were grown under moderate supply of nitrogen [N], and either high phosphate [P] (A-C,G,H), or low [P] (D-F,I-L). *Hyaloscypha hepaticicola* was used in all inoculation experiments. Different characters indicate differences (*P* < 0.05) between treatments, based on a Welch’s (double-tailed) *t*-test (C, F), or based on a Kruskal-Wallis test with post-hoc Dunn’s test (M). **(N, O)** Representative images of *H. hepaticicola* grown under moderate [N] supply, and high [P] (N), or low [P] (O).

Given that nutrient availability regulates the establishment of AMS in angiosperms and thalloid liverworts, promoting a symbiotic permissive state^6,23,24^, we hypothesized that P and N availability might regulate the establishment of ErM in *C. fissa*. We conducted inoculation experiments under a range of P supply that had no detrimental impact on the growth of *H. hepaticicola* cultures (Fig. S1). We applied these nutrient conditions to mock- or *H. hepaticicola*-inoculated *C. fissa* plants. Under conditions of low P supply, *C. fissa* inoculated with *H. hepaticicola* produced more biomass than mock-inoculated plants (Fig. 1D-F, and Fig. S1). This is in contrast with the minimal, or absence of, growth differences observed under high P supply (Fig. 1A-C, Fig. S1). Reducing N supply under high P led to increased mycorrhizal growth response (Fig. S1), indicating an interaction between the two nutrients, as in the case of AMS^25^ Similar to the growth response, low P supply allowed the development of intracellular hyphae in swollen rhizoid tips, while high P abolished colonization (Fig. 1G-M, Fig. S1).

In addition to AMS, the nitrogen-fixing root nodule endosymbiosis formed by few angiosperm species and nitrogen-fixing bacteria, is also regulated by nutrient availability^26^. Here we provide experimental evidence that nutrient availability defines symbiotic permissiveness in a third intracellular symbiosis. We propose that the regulation of colonization by the plant nutrient status is a key driver in maintaining mutualism irrespective of the symbiont type.

### Jungermanniales genomes

In order to provide genomic resources for Jungermanniopsida to further study ErM, we produced and annotated genome assemblies for *Calypogeia fissa* and *Solenostoma infuscum* (previously referred to as *Jungermannia infusca*). The *S. infuscum* genome was sequenced using the Illumina technology, yielding 248 million of paired-end reads assembled into 4,616 scaffolds (from 949bp to 1,4Mb, N50=244Kb) (Table S1). The *C. fissa* genome was sequenced using the PacBio long-read technology, resulting in an assembly comprising 82 scaffolds (from 46Kb to 66Mb, N50=16Mb, Table S1). As previously suggested by phylogenetic analyses conducted on transcriptome assemblies, the *C. fissa* and *S. infuscum* genomes both contain all four CSP genes *SYMRK, CCaMK, CYCLOPS* and *EPP1* (Fig. S2-5).

Whereas the estimated size of the *S. infuscum* genome is *ca*. 400 Mb, the estimated genome size of *C. fissa* is *ca*. 800 Mb. Genome assembly completeness analyses indicated 95.3% and 96% completion for *S. infuscum* and *C. fissa*, respectively, with 94.6% of the annotated genes duplicated in *C. fissa* (Table S1). Gene structural annotation predicted 14,494 and 38,150 protein-coding genes in the genomes of *S. infuscum* and *C. fissa* respectively (Table S1). Overall, the genome assembly statistics supported the hypothesis that *C. fissa* underwent whole-genome duplication (WGD), as previously proposed^27^ (Fig. 2A,B). This hypothesis is further supported by the distribution of synonymous substitutions per site (Ks) between pairs of paralogs in *C. fissa*, with a single peak near the origin (Ks = 0.05), which is indicative of a recent large-scale duplication event (Fig. 2A). Likewise, an intragenomic synteny analysis recovered 137 collinear blocks comprising 7,087 gene pairs (*i*.*e*., syntelogs) representing 37.2% of the *C. fissa* genome (Fig. S6), with nearly all of these corresponding to the peak near the origin (Ks = 0.05) (Fig. 2A). In addition to paralogous Ks, orthologous Ks was estimated for *C. fissa* and *S. infuscum*. The resulting Ks plot of orthologous genes exhibited a peak around Ks = 0.35, consistent with a scenario of recent WGD in *C. fissa* (Fig. S6) after its divergence from *S. infuscum*. Intergenomic comparison of synteny between *C. fissa* and the distantly related species *M. polymorpha* identified relatively few collinear gene pairs between them, roughly 10% of genes in each species, likely owing to the species’ deep phylogenetic divergence (Fig. S6). However, a clear 2:1 pattern of syntenic depth was observed among collinear genes (Fig. 2C), consistent with the apparent absence of WGD events in *M. polymorpha* (Fig. S6). The species tree inferred from orthologous genes recovered *C. fissa* and *S. infuscum* forming a clade (estimated divergence at 69.8 Ma, 95% HPD = 32 - 122 Ma) that is sister to the Marchantiopsida clade of complex thalloid liverworts (293.7 Ma, 95% HPD = 217 - 370 Ma) (Fig. 2D).

**Figure 2.**
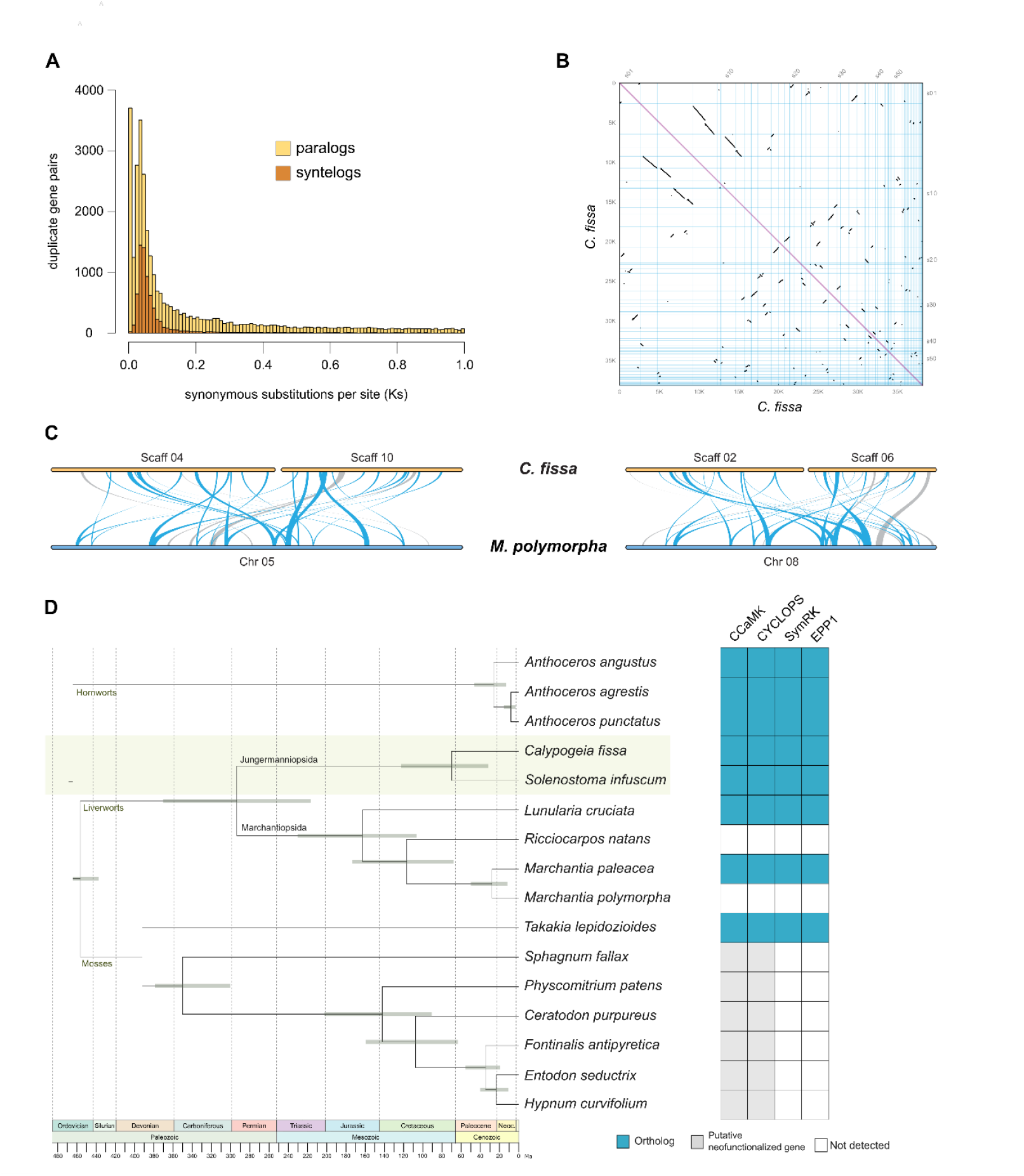
A recent whole-genome duplication in the leafy liverwort *Calypogeia fissa*. **(A)** Distribution of synonymous substitutions per site (Ks) between paralog pairs and syntelogs in the *C. fissa* genome. **(B)** Intragenomic synteny analysis identifies conserved collinear gene arrangements within the *C. fissa* genome, revealing 137 collinear blocks comprising 7,087 syntelogs and covering 37.2% of the *C. fissa* genome. **(C)** Synteny between *C. fissa* and *Marchantia polymorpha*. **(D)** Time-calibrated phylogeny of bryophyte species. Node bars represent the 95% highest posterior density (HPD) intervals. The root of the tree and the crown node of mosses were fixed to the mean ages from ^2105^ (see Methods). Presence of selected symbiosis-related genes is shown on the right panel.

Here, we have generated two genome assemblies for the Jungermanniales which are distantly related to the model Marchantiopsida liverworts.

### Induction of an ancestral gene module for intracellular symbioses

Availability of its genome assembly allowed us to conduct bulk RNA sequencing on mock-treated or *H. hepaticicola*-inoculated *C. fissa* plants, to test for the occurrence of a transcriptomic rewiring, as observed in other endosymbiosis^6,28–30^. For this, plants were grown in a permissive condition and harvested eight weeks after inoculation (Table S2). Differentially expressed genes were computed, identifying 787/743 genes up-/down-regulated (logFC threshold of |1|) in inoculated plants compared to the mock treatment in this condition permissive for ErM (Figure 5A, Supplementary Dataset S1). We hypothesized that comparing the transcriptomic response of *C. fissa* to the inoculation with *H. hepaticicola* in permissive and non-permissive symbiotic conditions would reveal intracellular-specific responses (Fig. 5A-B). To do so, we conducted RNA sequencing experiments with mock- or *H. hepaticicola*-inoculated *C. fissa* plants that were grown under a restrictive nutrient regime with high P supply for eight weeks. In this non-permissive condition, where no intracellular colonization was detected (Fig. 3A-C, 3G-H, and 3M), we computed 566/27 genes up/downregulated in the inoculated compared to mock-inoculated plants at a logFC of |1| (Fig. 5B, Supplementary Dataset S2). We compared the list of DEG in the permissive versus non-permissive conditions to identify the genes and processes associated with intracellular symbiosis during ErM, and conducted functional gene ontology enrichment in each condition (Fig. 5A-B, Tables S3-6). These comparisons revealed 82/1 genes up/downregulated in response to *H. hepaticicola* that were specifically responsive during the ErM-permissive state. Among the upregulated genes in permissive condition, we observed two highly enriched gene ontology categories, one encompassing a large number of enzymes associated with cell-wall remodeling and the other with nutrient transport (Fig. 5A-B, Table S3). Both categories might be involved in the reciprocal exchange of resources – carbon for nutrients – between *C. fissa* and the ErM fungus and for building the symbiotic interface.

**Figure 3.**
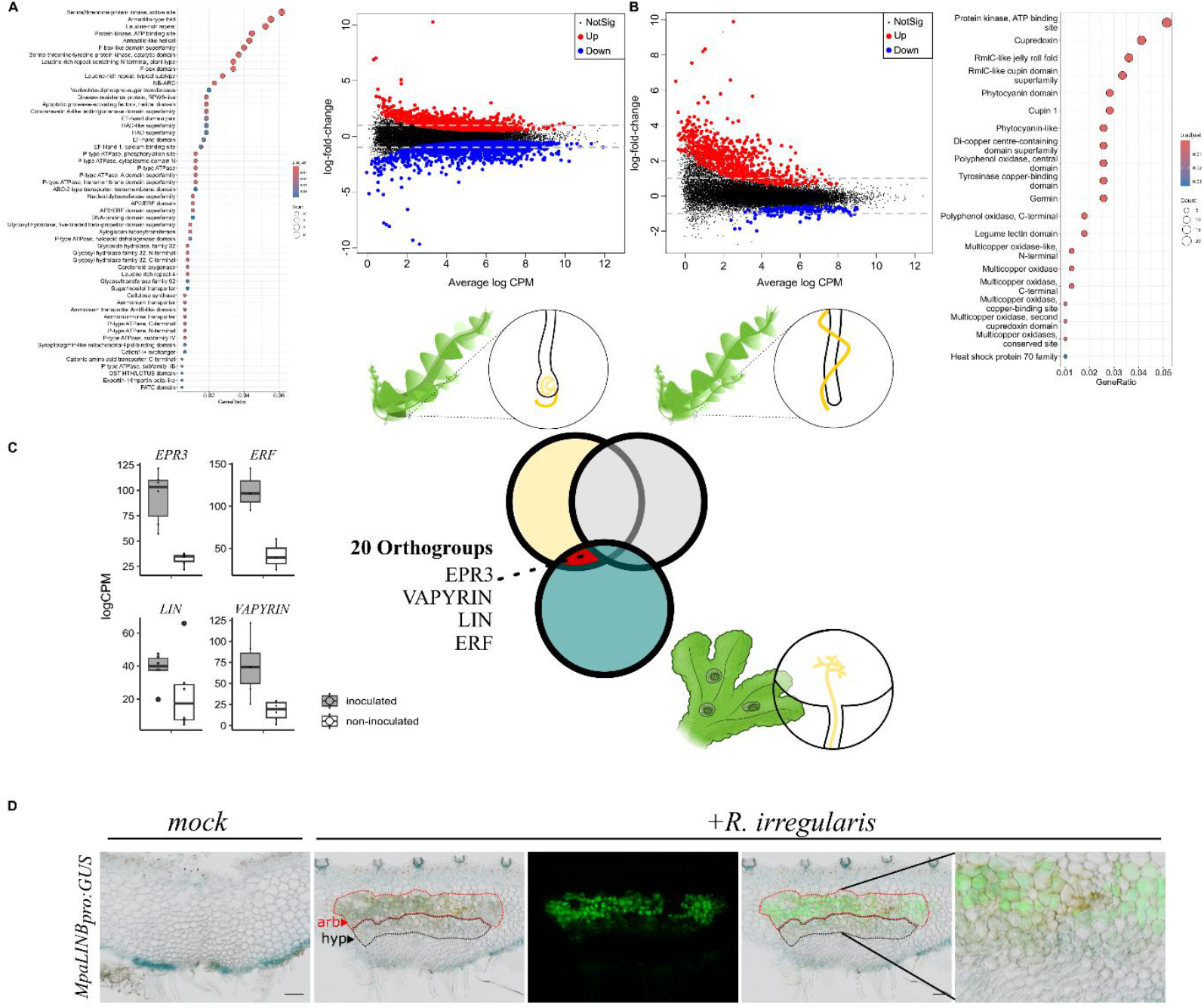
Transcriptional reprogramming during intracellular symbiosis. **(A)** Differential gene expression analysis comparing mock-treated and *Hyaloscypha hepaticicola*-inoculated *Calypogeia fissa* plants under ericoid mycorrhiza-permissive conditions. A total of 787 genes were upregulated, and 743 genes were downregulated in inoculated plants relative to mock-treated controls. **(B)** Differential gene expression analysis under ericoid mycorrhiza-permissive restrictive conditions, with 566/27 genes up/downregulated in inoculated plants compared to mock-treated controls. **(C)** Comparative transcriptomic analysis of ericoid mycorrhiza-permissive and restrictive conditions alongside arbuscular mycorrhiza-permissive conditions. The orange-overlapping area in the Venn diagram depicts the conserved expressed gene cluster that was found to be upregulated during both ericoid and arbuscular mycorrhizae. **(D)** Expression bar plots displaying expression of *C. fissa EPR3, ERF, LIN* and *VAPYRIN*. **(E)** Promoter activity of the *Marchantia paleacea LINb* gene monitored using *MpaLINBpro:GUS* at 6 weeks post inoculation. From left to right, mock inoculated (bright field), *R*.*irregularis-* inoculated (bright field), *R*.*irregularis-*inoculated (WGA-Alexafluor 488), *R*.*irregularis-* inoculated (merged), *R*.*irregularis-*inoculated (merged magnified). scale bar= 100 µM. Arrowheads indicate the the hyphae-containing area (hyp, black-dotted line), and the arbuscule area (arb, red-dotted line).

Next, we hypothesized that, as illustrated by the phylogentic pattern of the CSP genes (Fig. 2, Fig. S3-8)^5^, intracellular symbiosis might rely on shared gene modules, irrespective of the symbiont type and plant lineage. To test this hypothesis, we cross-referenced the specific transcriptomic signature of *C. fissa* in response to *H. hepaticicola* under the ErM-permissive condition to the transcriptomic response of the thalloid liverwort *Marchantia paleacea* colonized by the AMS-forming fungus *Rhizophagus irregularis*^6^ (see Materials & Methods). This analysis identified a total of 20 Orthogroups in *C. fissa* upregulated during ErM that had ortholog genes upregulated during AMS in *M. paleacea*. Among them, we detected the LysM-RLK receptor-encoding gene *EPR3* (Calfis_g18879 & Calfis_g34255), and the two genes, *VAPYRIN* (Calfis_g4017 & Calfis_g10154) and *LIN* (Calfis_g31157 & Calfis_g22484), encoding required proteins in angiosperms for the development of the intracellular symbiosis-specific structure called the infectosome (Table S7) ^31–35^. This suggests that the infectosome might be an ancestral feature induced during intracellular symbiosis in plants. To experimentally test this hypothesis, we used the genetically tractable liverwort *M. paleacea. M. paleacea* transformed with *MpaLINb*_*pro*_*:GUS* constructs were mock- or *R. irregularis-*inoculated and the GUS expression pattern was monitored 6 weeks later. In mock-inoculated conditions, lines transformed with the *MpaLINb*_*pro*_*:GUS* construct showed faint GUS activity in the ventral epidermis (Fig. 3). This basal expression was maintained in the plants colonized by *R. irregularis*. However, colonization by *R. irregularis* triggered and additional expression pattern in the area of the parenchyma hosting the fungus (Fig. 3).

Altogether, these RNAseq data across species and the promoter:GUS lines demonstrate that the intracellular symbionts-mediated local induction of the infectosome defines an ancestral symbiotic feature in land plants.

### ERS is essential for intracellular symbioses

To determine the mechanisms governing the induction of the infectosome, we searched for transcription factors in the 20 Orthogroups found up-regulated during both ErM and AMS and identified one GRAS (GRAS17) and one containing ERF transcription factors (Table S7). Although other GRAS proteins are involved in symbiotic processes^36–38^, their ability to directly bind DNA remains debatable. Thus, we decided to focus on the ERF transcription factors. This Orthogroup contains one gene in *C. fissa* (Calfis_g17180) and two paralogs in *M. paleacea* (Marpal_utg000001g0000321, Marpal_utg000040g0076701) which are up-regulated during AMS (Table S7). Promoter GUS fusions revealed basal level of expression for both *M. paleacea* genes in the thallus and the rhizoids in non-inoculated condition (Fig. 4). Upon colonization by *R. irregularis*, an additional expression domain was observed in the inner parenchyma hosting the AM fungus, mirroring the *MpaLINb*_*pro*_*:GUS* pattern (Fig 4). To test for the putative symbiotic function of this ERF Orthogroup in intracellular-symbiosis, we generated two single and two double mutants by CRISPR/Cas9 in *M. paleacea* (Fig. S9). The mutants and empty-vector control lines were inoculated with *R. irregularis* and harvested after 6 weeks. Thalli from the control showed accumulation of the AMS-induced pigment (Fig. 4), and displayed intracellular hyphae and arbuscules (Fig. 4). By contrast, not a single thallus of the two independent double mutant lines showed the accumulation of the pigment, or any colonization by *R. irregularis* (Fig. 4), mirroring the phenotypes of mutants in the CSP genes^18,19^. Given the complete lack of colonization of the double mutants, we named these ERFs *ERF Required for Symbiosis 1* and *2* (*ERS1* Marpal_utg000001g0000321 and *ERS2* Marpal_utg000040g0076701). Single mutants of one of the two paralogs, *MpaERS2*, displayed reduced colonization in three out of four independent experiments (Fig. S10), indicative of genetic redundancy between the two genes. To define the potential link of the ERS transcription factors with intracellular symbioses in other clades, we conducted a phylogenetic analysis on 127 plant species (Table S8). This analysis revealed 3 ERS clades. The first one is specific to non-flowering plants, including *MpaERS1* and *MpaERS2*, as well as the *C. fissa* ERS upregulated during ErM (Fig. 4). The other two clades likely originate from a duplication at the base of the angiosperms. One of these clades includes three genes from the legume *M. truncatula* – *ERN1, ERN2* and *ERN3* – which are known for their redundant symbiotic role during establishment of the nitrogen-fixing root nodule symbiosis in that species^39^ (Fig. 4). The second angiosperm clade encompasses three other *M. truncatula* genes including *ERM1. M. truncatula erm1* mutants show a quantitative defect in AM symbiosis^39^. We cross-referenced this phylogenetic data with transcriptomic data comparing mock-inoculated and symbiont-inoculated (root nodule or AM symbioses) plants available for 13 of the species represented on the tree. All tested species showed at least one ERS homolog up-regulated in response to AMS and/or RNS (Fig. 4, Table S9). Given this conserved symbiotic induction, and the phenotype of the *M. paleacea ers1ers2* mutant, we reasoned that ERS might regulate the expression of the infectosome. In angiosperm, Medicago ERN1 regulates gene expression via binding to the GCCGGC motif^40^. To determine whether this link might be conserved across land plants, we scanned the *VAPYRIN*_*pro*_ and *LIN*_*pro*_ sequences across land plant species for the presence of this motif, and detected it in, respectively, 106 out of 168 promoters (∼63%) and 147 out of 220 promoters (67%), indicating the conservation of the ERS – Infectosome logics between angiosperm and bryophytes (Table S10). To further test this link, we conducted *trans*-activation assays in *Nicotiana benthamiana* leaves with *MpaERS1* or *MpaERS2* on the *MpaLINb*_*pro*_. While the *MpaLINb*_*pro*_ did not show any basal expression (Fig. 4), co-expression with either of the two *MpaERS* led to its strong trans-activation (Fig. 4). Collectively, these data define *ERS* as a clade of transcription factors which have regulated the symbiotic induction of the infectosome across symbiotic types and plants lineages for the last 450 million years.

**Figure 4.**
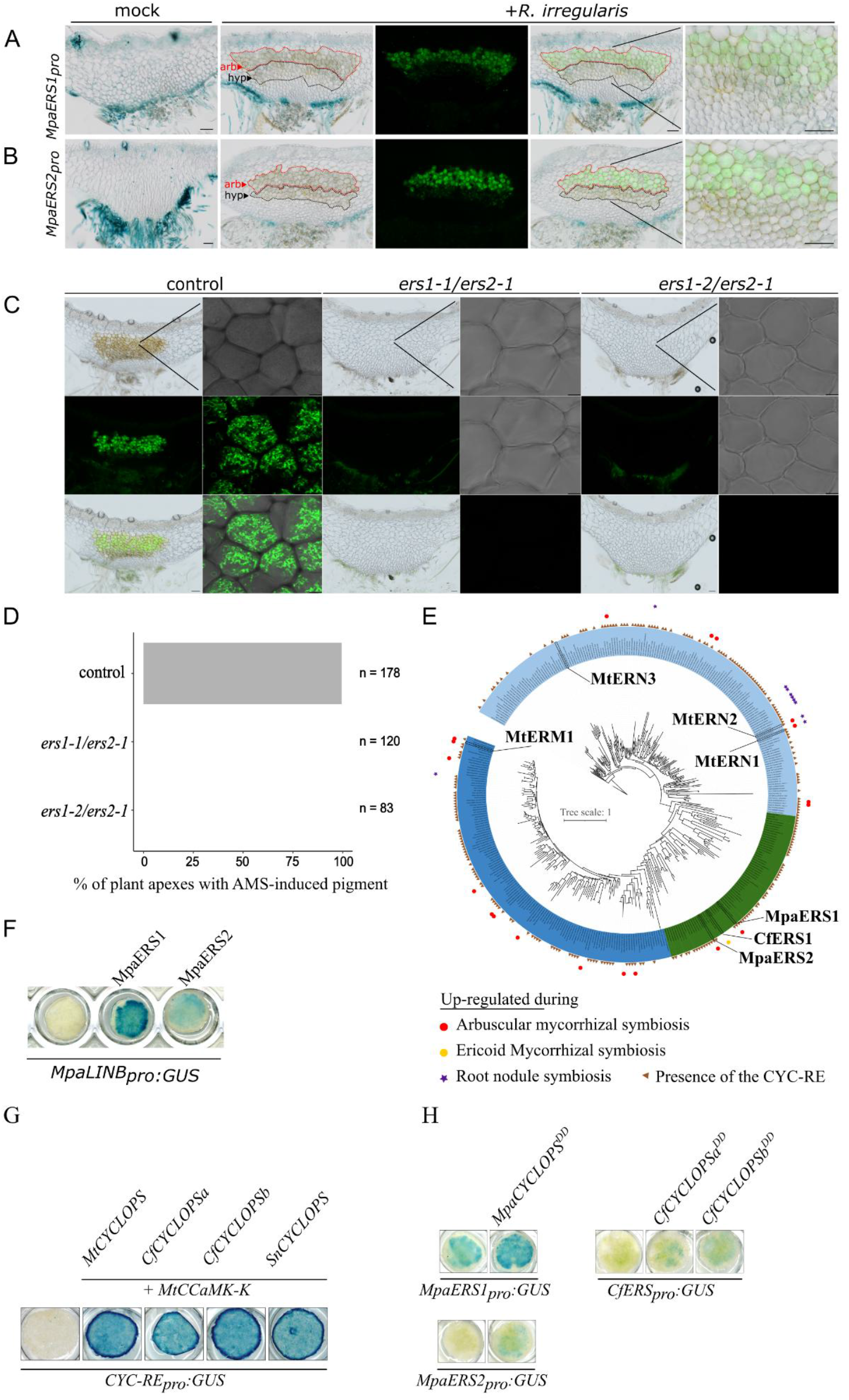
ERS bridge the CSP and the infectosome. **(A-B)** Promoter activity of the *Marchantia paleacea ERS1* and *ERS2* genes monitored using *MpaERS1pro:GUS* and *MpaERS2pro:GUS* at 6 weeks posr inoculation. From left to right, mock inoculated (bright field), *R*.*irregularis-*inoculated (bright field), *R*.*irregularis-*inoculated (WGA-Alexafluor 488), *R*.*irregularis-*inoculated (merged), *R*.*irregularis-*inoculated (merged magnified). scale bar= 100 µM. Arrowheads indicate the the hyphae-containing area (hyp, black-dotted line), and the arbuscule area (arb, red-dotted line). **(C)** Representative images of control plants transformed with an empty vector and *Mpaers* double mutants inoculated with *R. irregularis* at 6 weeks post inoculation. Green channel: WGA-Alexafluor 488 staining of fungal structures. Scale bar 100 µM or 10 µM. **(D)** Scoring of plant apexes with AMS-induced pigment at 6 weeks post inoculation with *R. irregularis* in control plants transformed with an empty vector and *Mpaers* double mutants. Combined results of 4 independent experiments are shown. **(E)** Reconstructed phylogeny of ERS orthologs across angiosperms (2 clades in blue) and bryopythes (clade in green), with the *CYC-RE* presence and up-regulated genes during different symbioses mapped on it. **(F)** *Nicotiana benthamiana* leaves expressing *MpaLINBpro:GUS* alone or together with *MpaERS1* or *MpaERS2*. **(G)** *N. benthamiana* leaves expressing *CYC-RE*_*pro*_*:GUS* alone or together with *MtCCAMK-K* and *CYCLOPS* from *Medicago truncatula* (*MtCYCLOPS*), or *Calypogeia fissa* (*CsCYCLOPSa and b*) or *Scapania nemorea* (*SnCYCLOPS*). **(H)** *N. benthamiana* leaves expressing *ERS*_pro_:*GUS* fusions from *Marchantia paleacea (MpaERS1*_*pro*_*:GUS, MpaERS2*_*pro*_*:GUS) and C. fissa* (*CsERS*_*pro*_*:GUS*) alone or together with constructs mimicking CYCLOPS phosphorylation (CYCLOPS^DD^) from *M. paleacea* (*MpaCYCLOPS*^DD^), or *Calypogeia fissa* (*CsCYCLOPSa*^*DD*^ and *CsCYCLOPSb*^*DD*^). GUS staining was performed at 48HPI and is shown in blue.

### Regulation of ERS by the CSP across plant symbioses

The CSP is a constitutively expressed pathway whose activation, mediated by the perception of symbiont-derived signals, leads to the phosphorylation of the transcription factor CYCLOPS by the kinase CCaMK. The CSP is phylogenetically linked with intracellular symbiosis^5^ (Fig. 2). Our data indicate that the ERS - infectosome module is induced by diverse symbionts across plant lineages and allows – at the cellular level – the actual intracellular colonization to occur. We reasoned that the direct regulation of ERS by phosphorylated CYCLOPS would integrate these two seemingly independent pathways, connecting symbiont perception to the transcriptional induction of the infection program. In angiosperms, this theory is supported in the context of the derived nitrogen-fixing root nodule symbiosis with the direct regulation of *ERN1* by CYCLOPS via a *cis-*regulatory element (CYC-RE) in the *ERN1*_*pro*_^41^ To determine whether this logic was conserved in Jungermanniales, we first conducted transactivation assays with the *CYCLOPS* ortholog genes of *C. fissa* (*CfCYCLOPSa* and *CfCYCLOPSb*) and another leafy liverwort, *Scapania nemorea* (*SnCYCLOPS*). As a control, we used *CYCLOPS* from *Medicago truncatula*^18^. We expressed either *CfCYCLOPS* or *SnCYCLOPS* in combination with an autoactive version of CCaMK (MtCCaMK-Kin)^42^ and the reporter construct *CYC-RE*_*pro*_*:GUS*^18,43^ in *N. benthamiana* leaves. As expected, no GUS activity was detected in *N. benthamiana* leaves transformed with *CYC-RE*_*pro*_*:GUS* alone (Fig. 4G). Strong GUS activity was observed upon co-transformation of the *MtCCaMK-Kin* (Fig. 4G) construct with either *CfCYCLOPS* (a and b), *SnCYCLOPS*, or with the positive *MtCYCLOPS* control construct. In addition, constructs mimicking CYCLOPS phosphorylation (*CYCLOPS*^*DD*44^, *CfCYCLOPSa*^*DD*^, *CfCYCLOPSb*^*DD*^ and *MpaCYCLOPS*^*DD*^, induced the *CYC-RE*_*pro*_*:GUS* (Fig. S11).

Using an untargeted promoter analysis, we found a highly conserved motif which corresponds to the CYC-RE in the promoters of the *ERS* homologs across land plants, further supporting the link between the CSP and ERS (Fig. 4, fig. S12). We evaluated whether the activation of *CYC-RE*_*pro*_*:GUS* by various CYCLOPS translated to the ERS promoters themselves. As for the *CYC-RE*_*pro*_*:GUS* above, we conducted transactivation assays using constructs mimicking CYCLOPS phosphorylation (CYCLOPS^DD^) from *M. paleacea* (*MpaCYCLOPS*^*DD*^) and *C. fissa* (*CsCYCLOPSa*^*DD*^, *CsCYCLOPSb*^*DD*^) (Fig. 4H) combined with either *MpaERS1*_*pro*_*:GUS, MpaERS2*_*pro*_*:GUS or CfERS*_*pro*_*:GUS*. The *MpaERS1*_*pro*_*:GUS* displayed basal expression, while the other two ERS promoters alone did not display GUS signals (Fig. 4H). Co-expression with any of the *CYCLOPS*^*DD*^ constructs from *M. paleacea* or *C. fissa* led to the induction of both *CfERS*_*pro*_*:GUS* and *MpaERS2*_*pro*_*:GUS* (Fig. 4), confirming the link between the CSP and the ERS.

Altogether, these phylogenetic and experimental data demonstrate that transcription factors of the ERS clade have been acting as a transcriptional link between the CSP and the infectosome since the most recent common ancestor of the land plants. Diversification of symbiotic options has been associated with the convergent recruitment of the CSP – ERS – Infectosome programme. Conceptually, the diversification of receptors upstream the CSP may be sufficient for this recruitment. The discovery that a conserved infectosome core module underlies the intracellular accommodation of completely distinct microbial endophytes in alternative symbiotic types, and the identification of a single clade of transcription factors associated with infectosome induction, provides valuable insights for guiding efforts to develop synthetic intracellular symbioses in land plants.

## Acknowledgments

We are grateful to Silvia Perotto (University of Turin) for kindly providing the *Hyaloscypha hepaticicola* (syn. *Rhizoscyphus ericae* UAMH 6735) strain used in this study. We thank the genotoul bioinformatics platform Toulouse Occitanie (Bioinfo Genotoul, https://doi.org/10.15454/1.5572369328961167E12), the NIG supercomputer at ROIS National Institute of Genetics, and the Data Integration and Analysis Facility at the National Institute for Basic Biology for providing computing resources. We acknowledge the TRI-FRAIB imaging facility, member of the national infrastructure France-BioImaging supported by the French National Research Agency (ANR-10-INBS-04). Celophane was provided by Futamura Chemical, Co., LTD. We thank Dr. Elizabeth Jamet (LRSV, CNRS) for kindly allowing us to use the camera for phenotyping. We are grateful to Dr. M. Shimamura for letting us know the change in taxonomy.

## Author contributions

Conceptualization: PMD, TV, KM, LCa, JK ; Methodology: ALR, CC, CLe, CLi, LC, LCh, MEB, OT, SA, TP, TV, KM, CRL, DW ; Investigation: CLi, LCa, LCh, MEB, SA,TP, TV, KM, DW, LF ; Visualization: LCa, MEB, SA, TV, KM, DW, SA ; Funding acquisition: PMD, KM, MEB ; Project administration: PMD ; Supervision: PMD, TV, KM, JK, FWL ; Writing – original draft: LCa,PMD, TV, KM, JK ; Writing – review & editing: all authors

## Data and materials availability

The *Solenostoma infuscum* genome assembly with annotation and raw reads are available on INSDC under the BioProject accession PRJDB19613. The *Calypogeia fissa* genome assembly and annotation is availablee on ENA-EMBL, with the reference number PRJEB86056. All DEG results, functional annotation, orthogroups and template scripts have been deposited in the repository 10.57745/MD77E8. An interactive Genome Browser can be found and queried here: https://bbric-pipelines.toulouse.inra.fr/myGenomeBrowser?browse=1&portalname=Calypogeia_fissa&owner=jean.keller@cnrs.fr&key=DKW4edLL. The raw RNAseq read of *Calypogeia fissa* are available on the SRA database, with the reference number PRJNA1186220. All read counts, orthogroups, differentially expressed genes and functional annotation of *C. fissa* can be accessed at: https://doi.org/10.57745/MD77E8.

## Supplementary Materials

Figure S1-S12

Tables S1 to S12

References (*45–105*)

