## Supplementary material for "Symbiotic diversification relies on an ancestral gene network in plants": Supplementary_materials_Castanedo et al.pdf

Materials and Methods
Figs. S1 to S12
Tables S1 to S2

### Materials and Methods

Preparation of high molecular weight (HMW) DNA for sequencing

*C. fissa* genomic DNA was isolated from whole plants using QIAGEN Genomic-tips 500/G kit (Cat No./ID: 10262) following the tissue protocol extraction. Briefly, 15g of plant material were frozen and ground in liquid nitrogen with mortar and pestle. After 1h of lysis at 50°C and one centrifugation step, the DNA was immobilized on the column. After several washing steps, DNA was eluted from the column, then desalted and concentrated by alcohol precipitation. The DNA was resuspended in EB buffer. DNA quality and quantity were assessed respectively using the Nanodrop-one spectrophotometer (Thermo Scientific) and the Qbit 3 Fluorometer using the Qbit dsDNA BR assay (Invitrogen). The size of the DNA was assessed using the FemtoPulse system (Agilent, Santa Clara, CA, USA).

DNA extraction and short reads sequencing

Genomic DNA from *S. infuscum* was extracted with the CTAB method and purified using a Qiagen genomic Tip 20. 180-bp and 700-bp insert Illumina paired-end libraries were constructed with a TruSeq PCR-free Sample Prep kit from covaris S2 fragmented and size-fractionated DNA. A 350-bp insert library was constructed with a gel-free protocol. Multiple mate-pair libraries were constructed with a Nextera mate pair library preparation kit. At first, fragmentation was carried out with 4 μg of DNA and three fractions were recovered after gel electrophoresis. A second mate-pair library set was constructed starting 10 µg of DNA and seven fractions were recovered after CHEF electrophoresis (switch time 1-4 sec, 16 hr, 0.5X TBE, 1% Megabase Agarose). Total RNA was extracted from 6-day old tissue using Qiagen RNA mini kit with RLT buffer. RNA-seq libraries were constructed with a TruSeq Stranded mRNA Prep kit with standard polyA selection or rRNA depletion usingRibo-Zero Magnetic kit (Plant leaf). Sequencing of the libraries were carried out on Illumina sequencers HiSEQ2000/2500 and MiSeq at the National Institute for Basic Biology, Okazaki, Japan.

Preparation and sequencing of HiFi PacBio library

A HiFi SMRTbell® library was prepared using the SMRTbell® Template Prep Kit 2.0 from Pacific Biosciences, following the manufacturer's instructions. High-molecular-weight DNA was fragmented to around 20 kb, then treated to remove single-strand overhangs and repair damage. Overhang adapters were attached to both ends of the DNA to form a closed circular DNA molecule, with further clean-up done using the SMRTbell® Enzyme Clean-up Kit 2.0. The library's size and concentration were measured with the FemtoPulse and Qubit systems. Sequencing primer and DNA polymerase were added before loading onto a SMRT cell, and sequencing ran for 30 hours on the Sequel® II system at the University of Lausanne, Switzerland.

Whole genome assembly

The *C.fissa* genome was assembled using HiFiasm assembler (v16.1)[45]. Hifiasm is able to produce a primary assembly and alternative assembly (incomplete alternative assembly consisting of haplotigs under heterozygous regions. We then filtered the primary assembly from organelles and low quality contigs. Organelles were identified by mapping contigs on NCBI reference assemblies. The closest references (NC_043787.1 and MF401632.1) were found by MitoHiFi software[46]. We finally removed 1009 contigs from the primary assembly (mitochondrial and chloroplastic contigs identified, high covered (>100X), low covered (<1X) and high %GC (>50%) contigs). All the metrics correspond to the filtered primary assembly used for the next analysis. We obtained 1,396,370 corrected reads (N50 = 17.4kbp; Genome Coverage = 32X) assembled and filtered in 82 contigs (N50 = 15.8Mbp; L50 = 14 contigs; GC content = 42.94%) for a total assembly size of

763Mbp. Genome assembly statistics were computed using the 'gaas_fasta_statistics.pl' script from the GAAS v1.3.0 suite (https://github.com/NBISweden/GAAS).

To assess the completeness and quality of the genome, we used the Benchmarking Universal Single-Copy Orthologs (BUSCO)[47] tool with the viridiplantae database (n=425). We obtained a 93.7% complete BUSCO score on the assembly.

The *S. infuscum* genome was assembled from Illumina short reads data using the ALLPATHS-LG program R49548 with default parameters.

Genome annotation

The *C. fissa* genome was first softmasked using EarlGrey v4.1.5[48] with default parameters and the NCBI taxid 33090 (Viridiplantae) for the RepeatMasker search. The -d option was set to 'yes' to produce a softmasked version of the genome.

Gene prediction in the *C. fissa* genome was done using BRAKER3.0.7[49] in ETP mode with both RNASeq and protein hints. RNASeq hints were produced from the transcriptomic data generated in this study and described below. Briefly, each sample was first mapped against the *C. fissa* genome using HISAT2 v2.2.1[50] with the following parameters: --sensitive --no-discordant --no-mixed --dta --max-intronlen 5000. Then each mapping file was processed with Samtools v1.19[51] collate, fixmate (with the -m -c and -r options), sort and markdup. Finally, all mapping files were merged into a unique file with the merge and collate options of Samtools prior being processed as described for individual mappings. Both mapping and protein database files were used by BRAKER to annotate the genome using the following tools as contained in the BRAKER 3.0.7 Singularity container (available at: https://hub.docker.com/r/teambraker/braker3): miniprot[52], GeneMark-ETP[53], diamond[58,], Spaln2[59], StringTie2[56], GFF utilities[57], AUGUSTUS[58,59] and TSEBRA[60]. To maximize the completion of the prediction, BRAKER was run with compleasm[61] option set to "Viridiplantae".

BRAKER prediction was then processed to remove the alternative transcript using agat_sp_keep_longest_isoform.pl from the AGAT v1.4.0 toolkit[62] and coding-sequence as well proteins extracted from the genome using gffread v0.12.8 from GFF utilities package[57] with the -C and -V options enabled. Proteins were functionally annotated uysing InterproScan5 v64.96.0[63] with the options '-iprlookup' and 'goterms' enabled.

The *S. infuscum* genome has been annotated using RepeatModeler Version open-1.0.7, RepeatMasker version open-4.0.5 (NCBI/RMBLAST 2.2.27+), exonerate (2.2.0), bowtie2 (2.1.0), tophat2 (2.0.10), bowtie (1.0.0), and AUGUSTUS v. 3.0.2 using scripts available at https://github.com/tomoakin/Jin-annot. Arabidopsis (TAIR10) and *Physcomitrium patens* (*P.patens*.V6_filtered_cosmoss_proteins.fas) protein datasets were used for exonerate mapping. 224 loci were manually checked on Web Apollo[64] and converted to genbank format for training Augustus, *Marchantia polymorpha* proteins collected from GenBank as of 1 Feb, 2017 were additionally mapped to the reference with exonerate. RNA-seq data were remapped with hisat2, and the final run using with UTR=on (version 3.2.3).

Whole genome duplication analysis in *C. fissa*

Paralogous Ks was calculated using the wgd package v1.1.2[65]. Specifically, genes were clustered with the mcl command using an inflation index of 2.0 prior to estimating paralogous Ks with the ksd command using the --pairwise flag. Orthologous Ks was estimated between *C. fissa* and *J. infusca* in a similar manner using the mcl command using the --one_v_one and with the ksd flag under default settings.

Syntenic gene pairs were identified and plotted using the python version of MCSCAN as implemented in the jcvi comparative genomics toolkit v0.9.2[66]. First, a file containing CDS sequences was generated with gffread[57] using the -x flag and the annotation file was converted from gff3 to bed format using the 'jcvi.formats.gff bed' function for *C. fissa* and *M. polymorpha*. The resulting CDS and bed files were used as input 'jcvi.compara.catalog ortholog' function to identify and visualize synteny within and between species. A default C-score of 0.7 was used to filter low quality hits.

Plant and fungal material, growth conditions, inoculation assays, and nutrient supply experiment of *C. fissa*

*Calypogeia fissa* (L.) Raddi ssp. *fissa* was sourced from the Edimburgh botanical garden and maintained in polypropylene microboxes (OV80 + OVD80L#10G, Sac O2, Dutscher no. 017055, Issy-les-Moulineaux, FR) with 150 g of silica sand (Sibelco, Puel, no. EMB33284, Toulouse, FR). Microboxes were supplemented with 40 mL Long-Ashton solution (standard P, high N; see Supplementary Table 1) and incubated at 23 °C, 16-hour day/ 8-night at 20 °C cycle. For the time-series experiments, plants were watered at weeks 4 and 7 post-inoculation. Permissive and non-permissive nutrient solutions for ericoid mycorrhizae in *C. fissa* were selected by screening the effects of seven different recipes with varying phosphate and nitrogen concentrations, based on the Long Ashton nutrient solution[67]. The selection was made by assessing plant and fungal growth, as well as colonization levels of *C. fissa* rhizoid tips by *H. hepaticicola*. Recipes for the nutrient solutions used in this study can be found in Supplementary Table 1.

*Hyaloscypha hepaticicola* (syn. *Rhizoscyphus ericae* UMAH 7357) (D.J. Read) Zhuang & Korf was maintained via mycelial agar plug transfers on Czapek-Dox (C-Dox) agar plates prepared per Atlas[68]. Before *C. fissa* cultivation, three agar plugs were transferred to microboxes containing silica sand and nutrient solution and incubated in dark conditions at 17°C for two weeks. Subsequently, ten *C. fissa* stems were introduced into each microbox, which further developed into plant colonies. For nutrient supply studies on plates, *R. ericae* agar plugs were added to plates with a nutrient medium consisting of working solutions from Supplementary Table 1, supplemented with 3.5% sucrose and 1.5% BD Bacto™ Dehydrated Agar (Fisher Scientific, no. 214010). Plates were incubated as described above.

*Solenostoma infuscum* (Mitt.) Hentschel (*syn J. infusca*) has been maintained as axenic culture at Kumamoto University and provided by Prof. Hiroyoshi Takano. The plants were grown on KNOP agar plates covered with a sheet of cellophane (provided by Futamura Chemical, Co., LTD.) after cutting with a Polytron similar to *Physcomitrium patens* culture but shorter time of cutting than protonema culture.

Phenotyping of *C. fissa*

Imaging of microboxes and fungal plates was conducted using a commercially available Canon EOS 4000D digital single-lens reflex camera (18.0 megapixels). Weekly images of microboxes containing three *C. fissa* colonies were captured to monitor growth. Colony area was estimated using ImageJ software[69]. The scale was set using the known width of the microbox (12.5 cm), and color thresholds were adjusted to optimize hue (35–30 upper, 255 lower), saturation (51 upper, 255 lower), and brightness (0 upper, 255 lower) for each image. Particle analysis was then performed with parameters set to a size range of 10,000–Infinity $cm^2$, circularity of 0.00–1.00, and results displayed as outlines. Outputs, including the processed image, analyzed data, and measurement summary, were saved for further analysis. The colony area ($cm^2$) measured weekly is reported as a proxy for biomass change in this study.

Microscopy and quantification of colonization events of *C. fissa*

Ten independent stems from three independent microbox biological replicates from inoculated and non-inoculated *C. fissa* samples were prepared for microscopy using Wheat Germ Agglutinin (WGA-cf488a) and Congo Red to visualize chitinous structures and cell walls. Samples were fixed in 10% (v/v) at room temperature overnight, washed three times in water, and resuspended in 1X (v/v) phosphate-buffered saline (PBS), containing 1 µg/mL WGA (Dominique Dutscher, no. 462396, Issy-les-Moulineaux, FR), and incubated overnight at room temperature in the dark. For Congo Red staining, samples were incubated in 0.5% congo red aqueus solution (Rouge Congo pur RAL Diagnostics, LABELIANS, no. 3634100005, Nemours, FR) for 10 minutes just before confocal acquisitions. After staining, samples were rinsed twice in water to remove excess dye and mounted on microscope slides in water to be observed.

Microscopy was performed using both epifluorescence and confocal microscopy. Epifluorescence imaging was conducted using a Zeiss Axiozoom V16 equipped with objective 0.5x, GFP like filter set (Zeiss 38 He filter set, ex:470/40, em: 525/50) and RFP filter set (Zeiss 20 filter set, ex:546/12, em: 575-640) for WGA and Congo Red visualization, respectively. We also used Nikon Eclipse Ti wide field microscope with Nikon GFP filter set (ex: 472/30, em: 520/35). Confocal microscopy was carried out using a Leica SP8 CSU and objective 25x/0.95 water immersion. WGA was excited at 488nm and emission set between 500 and 545 nm. Congo Red was excited at 552nm and fluorescence was collected between 590 and 640 nm. Z-stack was performed with step size range between 0.57µm to 1µm.

Image analysis and processing were performed using ImageJ and LAS X Life Science Microscope Software, with identical settings applied across all samples to ensure consistency.

Manual counting of *C. fissa* swollen rhizoid tips was performed using a four-digit manual cell counter (4.3 × 3.5 mm with reset knob, Avantor VWR no. MOIN1494MW, Rosny-sous-Bois, FR) under a Zeiss Axio Zoom.V16 microscope. Results obtained from the Axio Zoom.V16 were independently validated using a Nikon Eclipse Ti microscope.

Harvesting of plants and RNA extraction of *C. fissa*

Whole *C. fissa* colonies were harvested during the morning on week 8 post-inoculation. Plants were pooled from three replicate microboxes (three independent colonies with hundreds of plant individuals per colony) per experiment, frozen in liquid nitrogen and stored in 50-mL screw-cap polypropylene tubes at −80°C until further processing. Frozen plant tissues were homogenized in 50-ml polypropylene tubes (Avantor Performance Materials GmbH, Griesheim, Germany, Catalog no. 734-0453) pre-chilled in liquid N2, each containing a commercial glass marble, using a vortex (Vortex-Genie 2, Scientific Industries, Inc., Bohemia, New York, USA) at maximal speed for 10 s, with three to four cycles of grinding and chilling in liquid N2. RNA was isolated from aliquots of 300 mg of frozen homogenized tissue using the Direct-zol RNA Microprep Kit following supplier's instructions (Zymo Research Europe GmbH, Catalog no. R2062, Freiburg, DE).

Estimation of differentially expressed genes of *C. fissa*

RNASeq raw reads from all described conditions were mapped against the genome of *C. fissa* and counted using the Nextflow v23.10.0[70] pipeline NF-CORE/RNASeq v3.14[71], the options star_salmon to align and quantify reads as well as '-nextseq 30 -length 50' as extra parameters of TrimGalore v0.6.7 (10.5281/zenodo.5127899) to remove reads with quality lower than 30 or a length lower than 50bp. Ribosomal RNA were also removed through the option "-remove_ribo" using SortMeRNA v4.3.4[72]. The pipeline was run under the GenoToul configuration (https://github.com/nf-core/configs/blob/master/docs/genotoul.md) and used the following softwares and languages: bedtools v2.30.0, R v4.0.3, v4.1.1 and v4.2.1[73], DESEQ2 v1.28.0[74]; dupradar v1.28.0[75], fastqc v 0.12.1 (bioinformatics.babraham.ac.uk/projects/fastqc/), fq v0.9.1 (https://github.com/stjude-rust-labs/fq), gffread v0.12.1[57], perl v5.26.2 (dev.perl.org/perl5/news/), python v3.9.5 (python.org/downloads/release/python-395/), rsem v1.3.1[76], STAR v2.7.10a[77], picard v3.0.0 (broadinstitute.github.io/picard), qualimap v2.3[78], rseqc v5.02[79], salmon v1.10.1[80], summarizedExperiment v1.24.0[81], samtools v1.16.1[51], stringtie v2.2.1[56], tximeta v1.12.0[82], UCSC v377 and v445 (https://github.com/ucscGenomeBrowser/kent).

For each condition and time point, differentially expressed genes (DEGs) were estimated independently with edgeR v4.2.1[83] in R v4.4.0 using the quasi-likelihood F-tests method and a FDR threshold of 0.05.

Enriched functions were determined using the R package clusterProfiler v4.12.3[84], the DEGs with a logFC threshold of |1| as foreground and genes subjected DEG analysis (*i.e.*, genes that were retained after removing the low counts) as background.

Re-estimation of differentially expressed genes under AMS and RNS

Data from different studies were used to re-estimate deregulated genes during AMS on 6 different species (see Table S12). Treatment of reads and estimation of DEGs were done with the exact same pipeline as described above for *C. fissa*. For RNS, data was retrieved from Libourel et al. 2023[85].

Orthogroups reconstruction

Orthogroups at the scale of Bryophytes were reconstructed using OrthoFinder v2.5.5[86] covering sequenced Bryophyte lineages (list of included species is indicated in the Table S8, column 'used_for_orthogroups_reconstruction'). Prior to orthogroup reconstruction, genome annotation of each species was processed as described for *C. fissa*: retention of longest isoforms with AGAT and then extraction of sequences with gffread and the "-C -V" options. OrthoFinder was then run with the "diamond_ultra_sens" and the msa options[87]. For all downstream orthogroup-based analyses presented in this work, the phylogenetic hierarchical orthogroups containing all species (N0) has been used.

Cross-referencing of transcriptomic data and orthogroups

Orthogroups were then filtered to retain the ones containing only up-regulated genes of *C. fissa* in the ErM-permissive condition, no differential expressed genes in the restrictive condition and at least one gene up-regulated in *M. paleacea* at 3, 5 or 8 weeks post-inoculation.

Ortholog inference of symbiosis-related genes

We used BLAST v2.15.0[88] to search for putative homologs in the genomes of selected symbiotic and nonsymbiotic species of land plants (see Table S8), with the protein sequences of *Marchantia paleacea* and *Medicago truncatula* as queries (*e-value*= $10^{-9}$). Sequences were aligned using MAFFT v7.505[89] with the automatic mode (--*auto*, --*maxiterate* 20), and alignments were subsequently trimmed using trimAl v1.4.1[90] to remove columns with more than 70% of gaps (-*gt* 0.3). An approximately maximum likelihood tree was then estimated using FastTree v2.1.11[91] with the gamma model (-*gamma*). Trees were inspected to identify the group of orthologous sequences containing the query, which was extracted, realigned using MUSCLE v5.1.0[92] and trimmed with trimAl to remove columns with more than 95% of gaps (-*gt* 0.05). A maximum-likelihood tree was then estimated with IQ-TREE v2.2.2.6[93] using the standard model selection (-*m TEST*), with branch support assessed using the SH-like approximate likelihood test and ultrafast bootstrap, both with 1000 replicates.

Promoter motif analyses across ERS, VAPYRIN, and LIN orthologs

Promoter sequences of 3kb upstream were extracted from each gene from each phylogeny. Sequences containing more than 10% unknown nucleotides ('N') were excluded from analyses. To identify conserve sequence motifs in the promoter sets, the program MEME v5.5.7[94] was used (with the following parameters: '-dna -mod zoops -minw 5 -maxw 30 -nmotifs 30'. To complement this, known cis-regulatory elements were searched in the promoter sequences using a through regular expressions.

For ERS orthologs, 368 promoter sequences (out of 380 after quality filtering) were analyzed and the first motif found was corresponding to the palindromic sequence of the *CYC-RE*. Motif pattern searches were done with the regular expression: 'CCA**TGG', and it was found among 235 promoters (~64%).

For both VAPYRIN and LIN genes, 168 and 212 promoters were respectively retained after filtering and the ERN known motif 'GCCGGC'[95]was screened in each of them.

Estimation of divergence times

Divergence times among bryophyte lineages were estimated using a set of nuclear orthologous genes extracted from the orthogroup data set (see above). Because no single-copy orthogroups were detected, we selected those with duplicated genes in only one species (total = 346 orthogroups), in which case only the longest of the duplicates was retained. Orthogroups containing putative pseudogenes or non-homologous sequences were then identified and removed from the data set. For that, nucleotide sequences were codon-aligned as described above, and phylogenetic trees were inferred using IQ-TREE with the standard model selection. We then used TreeShrink v1.3.9[96] to identify orthogroup trees containing outlier long branches with a false positive rate of 5% ($q$ = 0.05), and removed these orthogroups from the data set. Finally, we randomly selected 50 orthogroup alignments, which were concatenated and trimmed using trimAl to remove all positions containing gaps (-*gt* = 1; final alignment length = 60,594 bp). A time-calibrated phylogenetic tree was estimated using BEAST v2.7.4[97]. We used the GTR+G substitution model with four gamma categories, with the Yule model as speciation prior and an optimised relaxed clock[98]. We adopted here a secondary calibration strategy using divergence time estimates from published data[99], which represents, to our knowledge, the most comprehensive assessment to date of the fossil record of land plants for node calibration of molecular dating. We fixed the age of the root at 464.5 Ma, and the age of crown mosses at 392 Ma, both ages corresponding to the respective mean values from Harris et al.[23], and that were used here as the mean of a normal distribution with standard deviation of 0.001. To incorporate the current understanding of the phylogeny of bryophytes and reduce tree search space, we fixed the position of *Takakia lepidozioides* as sister to the other mosses[100] and the monophyly of the three main bryophyte lineages[99]. Two MCMC chains were run in parallel for 400,000,000 generations each, sampling every 10,000 generations. We monitored the runs using Tracer v1.7.1[101] to check for convergence and effective sample sizes > 200 for all parameters. We discarded trees sampled before the convergence of the runs (burn-in = 10%) and combined all the remaining trees into a maximum clade credibility tree, with the median ages summarized on the nodes.

*Nicotiana benthamiana* transactivation assays

The Golden Gate modular cloning system was used to prepare the plasmids as previously published[6,102,103]. Levels 0, 1, and 2 constructs used in these assays are listed in Table S11 and held for distribution in the ENSA project core collection (https://www.ensa.ac.uk/). All sequences were domesticated, synthesized, and cloned into pMS (GeneArt, Thermo Fisher Scientific, Waltham). *MtCCaMK-Kin* and *CYC-RE*$_{pro}$ sequences were taken from published data[18].

Constructs were introduced into *Agrobacterium tumefaciens* GV3101 and co-infiltrated into *N. benthamiana* leaves as described elsewhere[104]. For co-infiltration, equal volumes of each *A. tumefaciens* different cultures were adjusted to an $OD_{600}$ of 0.5 and mixed prior to infiltration. Each combination of constructs was infiltrated into two different leaves of three different plants. 48 hours *post* inoculation, leaf discs were harvested and GUS stained as previously described[18] with the following modifications: 0.1% triton was added to the GUS solution and leaf discs were first placed under vacuum for 20 min in the GUS solution before incubation at 37°C for 20 hours. Discs were rinsed with water and treated with ethanol 95% to remove the chlorophyll.

*Marchantia paleacea* stable transformation

The Golden Gate molecular cloning system was used to generate constructs for promoter-reporter assays and CRISPR-Cas9 mediated targeted mutagenesis as previously published[6,102,103]. Level 0,

1, and 2 constructs and guideRNA sequences are listed in Table S11.To generate transformants of *M. paleacea*, wildtype gemmae from axenic plants were grown on Gamborgs B5 half strength medium (G5768, Sigma, France) pH 5.7, 1.4% bacteriological agar (1330, Euromedex, France) for 4 to 5 weeks in a walk-in growth chamber at 20 °C, 16 h light/8 h dark cycle. For each construct, 15 to 25 gemmalings were blended for 15 s in a sterile 250 ml stainless steel bowl (Waring, USA) in 10 ml of 0M51C medium ($KNO_3$ 2 g/L, $NH_4NO_3$ 0.4 g/L, $MgSO_4$ $7H_2O$ 0.37 g/L, $CaCl_2$ $2H_2O$ 0.3 g/L, $KH_2PO_4$ 0.275 g/L, L-glutamine 0.3 g/L, casamino-acids 1 g/L, $Na_2$MoO4 $2H_2O$ 0.25 mg/L, $CuSO_4$ $5H_2O$ 0.025 mg/L, $CoCl_2$ $6H_2O$ 0.025 mg/L, $ZnSO_4$ 7H2O 2 mg/L, $MnSO_4$ $H_2O$ 10 mg/L, $H_3BO_3$ 3 mg/L, KI 0.75 mg/L, EDTA ferric sodium 36.7 mg/L, myo-inositol 100 mg/L, nicotinic acid 1 mg/L, pyridoxine HCl 1 mg/L, thiamine HCl 10 mg/L). The plant tissues were transferred to 250 ml erlenmeyer flasks containing 15 ml of 0M51C and kept at 20 °C under 16 h light/8 h dark cycle on a shaker at 200 rpm for 3 days. *Agrobacterium tumefaciens* GV3101 transformed with the respective constructs were grown to saturating $OD_{600nm}$ at 28 °C. *M. paleacea* blended tissues were inoculated with 100 µL of saturated Agrobacterium culture and supplemented with acetosyringone (100 µM final). After 3 days of co-culture at 20 °C under 16 h light/8 h dark cycle on a shaker at 200 rpm, the plant fragments were washed three times with water, and plated on Gamborgs B5 half strength solid medium containing 200 mg/L amoxycilin (Levmentin, Laboratoires Delbert, FR) and 10 mg/L Hygromycin (Duchefa Biochimie, France). Transformants were selected after 3-5 weeks at 20 °C under 16 h light/8 h dark cycle.

Marchantia paleacea mycorrhization tests and promoter-GUS reporter assays

Five *M. paleacea* plants per pot were grown in 7×7×8 cm pots on a zeolite substrate (50% fraction 1.0 to 2.5 mm, 50% fraction 0.5 to 1.0-mm, Symbiom) in a walk-in growth chamber at 20 °C under 16 h light/8 h dark cycle for 2 weeks. Each pot was then mock-inoculated or inoculated with approximately 1,000 sterile spores (250 spores per plant) of *Rhizophagus irregularis* DAOM 197198 provided by Agronutrition (Labège, France, https://www.agronutrition.com/en/contact-us/) and co-cultivated for another 6 weeks in the growth chamber and watered once a week with Long-Ashton solution containing 15 µM of phosphate. At 6 weeks post inoculation, the thalli were harvested. For promoter-GUS reporter assays, the plants were submerged in GUS buffer (phosphate buffer (0.1 M), EDTA (5 mM), $K_3Fe(CN)_6$ (0.5 mM), $K_4Fe(CN)_6$ (0.5 mM), X-Gluc (0.25 mg/ml, Euromedex, France), and $H_2O$) directly after harvesting and vacuum infiltrated for

20 min at 200-300 mbar in an exicator, followed by overnight incubation at 37 °C in the dark. The GUS-buffer was removed subsequently and the plants were washed once with water. All plants were cleared of chlorophyll using 100% ethanol for 24 h and then stored in an aqueous solution containing EDTA (0.5 mM) until further processing.

*Marchantia paleacea* phenotyping

Cleared *M. paleacea* thalli were scanned and scored as previously described[6] using the presence of black/purple pigment as an indicator for colonization. To confirm presence or lack of colonization the thalli were subsequently subjected to microscopic imaging. For this, thalli were embedded in 7% agarose and 100 µM or 110 µM transversal sections were prepared using a Leica vt1000s vibratome. The sections were incubated overnight in 10% or 12% KOH and washed 3 times with water. To visualize arbuscules, the sections were incubated in 1µg/ml WGA-Alexa Fluor 488 (Invitrogen, France) in PBS buffer overnight at 4 °C in the dark. Sections were imaged using a Nikon Ti Eclipse inverted microscope equipped with DS Ri2 camera and motorized XY stage. Large images of global sections were generated using the NISAR 4.3 scan large image module that allow multifield acquisition and images stitching. These images were acquired with 10×/0.3 dry objective (0.73 pixel size) in brightfield and in fluorescence for WGA-Alexa 488 staining using a GFP band pass filter set (ex: 472/30 nm, em:520/35 nm). Close-up images were acquired using a Leica SP8 TCSPC confocal microscope and LAS X software with a 25× water immersion objective (Fluotar VISIR 25×/0.95 WATER) at zoom 5× (0.182 µm pixel size). WGA-Alexa 488 staining of fungi was excited with the 488 nm laser line and fluorescence was recovered between 500 nm and 550 nm. Brightfield image was also performed and merged. Images were processed with ImageJ.

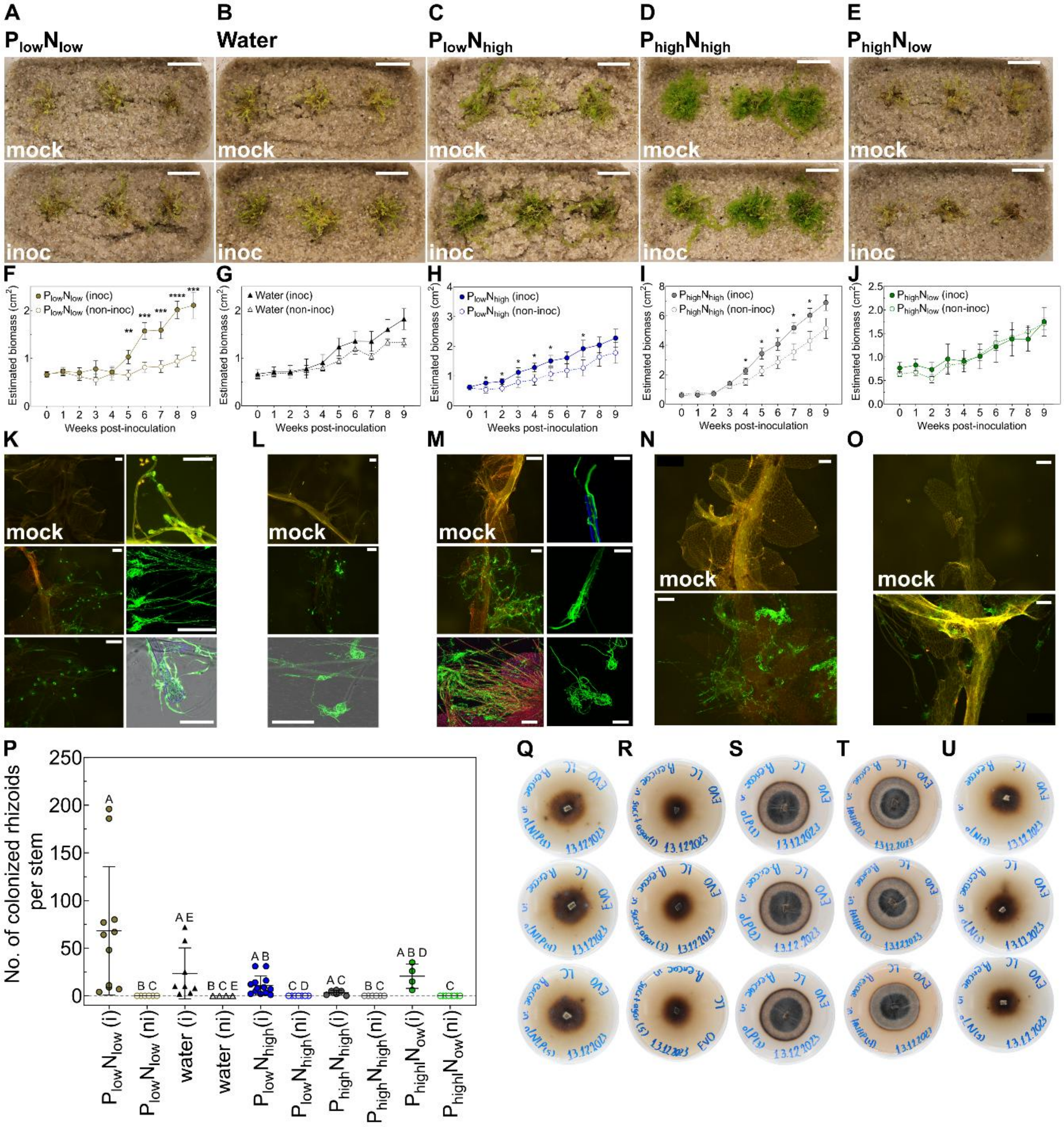

**Fig. S1. Establishment of a model to study ericoid mycorrhizae**. (A-J) Representative images (A-E), and estimated growth (F-J) of *C. fissa* plants in mock-inoculated (non-inoc) and inoculated (inoc) treatments. Scale bars: 2 cm. Data points in (F-J) indicate the mean ± SEM (n = 27) of technical replicates per time-point from one representative experiment of atotal of three independent experiments. (K-O) Representative fluorescence microscopy and confocal images of *C. fissa* stems harvested from non-incoc and inoc treatment. Stems were stained with WGA (green-fluorescent signal), and further stained with Congo red (blue- fluorescent signal). Scale bars: 200 µm, 75 µm, and 50 µm. (P) Total number of swollen and colonized rhizoid tips per stem in non-inoc (ni) and inoc (i) treatments. Data points indicate the mean ± SD (n = 4 to 13) of individual technical stem replicates from one representative experiment. Data is based on individual quantification of intracellular colonized rhizoids stained with WGA under epifluorescence and confocal microscopes. Different characters indicate statistically significant differences ($P < 0.05$) based on a Welch's (double-tailed) t-test (F-J), or based on a Kruskal-Wallis test with post-hoc Dunn's test (P). (Q-U) Representative images of *H. hepaticicola* grown under the indicated nutrient conditions in (A-E respectively) used for physiological characterization of C. fissa (F-J), or a basal medium consisting of water, 1.5% (w/v) sucrose, and 0.8% (w/v) Bacto agar (see Methods). Plates were incubated at 17°C in the darkness and photographed at week 23. All plants were photographed and harvested after 8 weeks of growth. Plants were grown under under the conditions indicated in (A-E). *H. hepaticicola* was used in all inoculation experiments. See Methods for details, Table S2 for nutrient recipes.

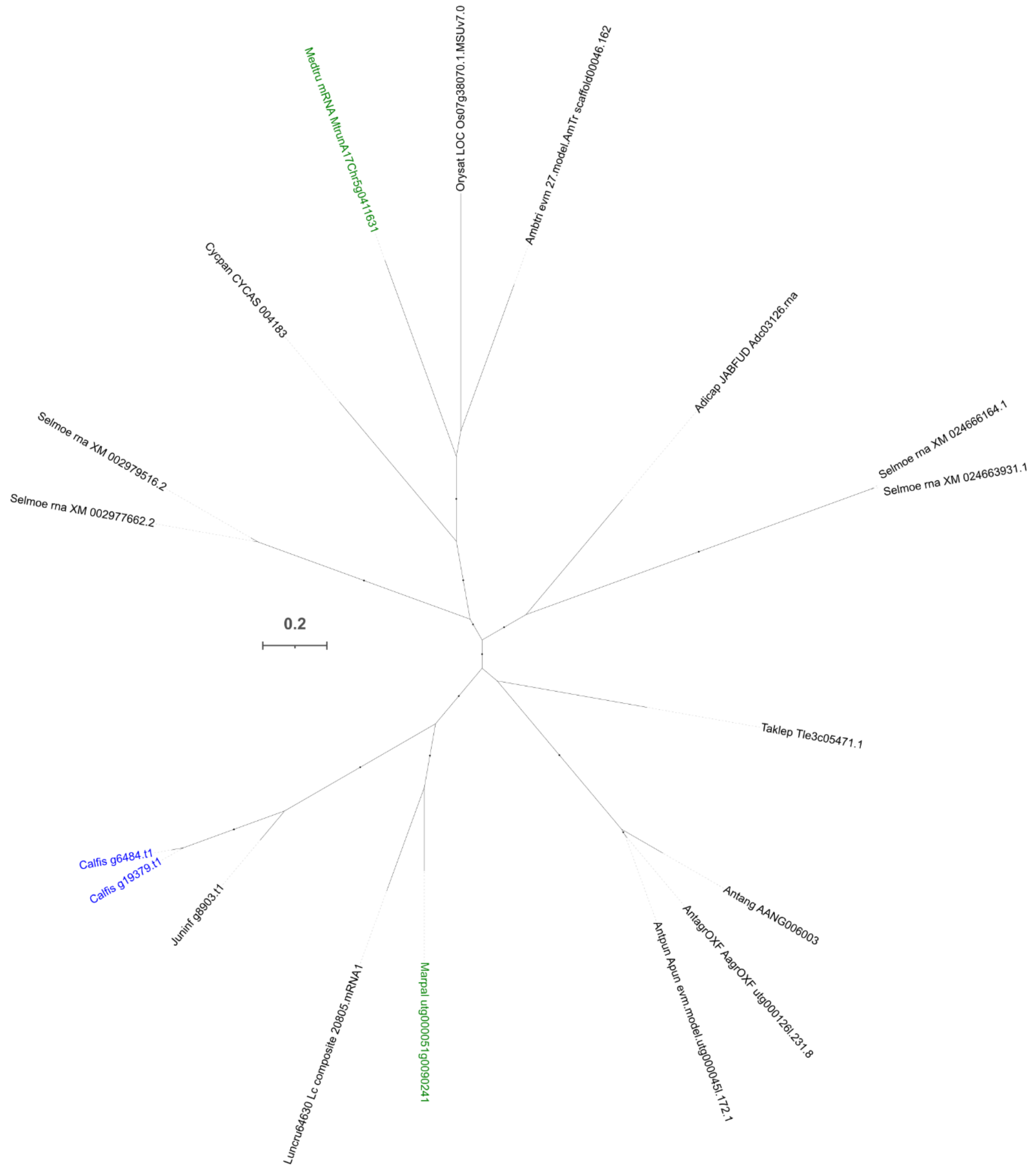


**Fig. S2. Phylogenetic tree of *SymRK*.** Maximum-likelihood tree estimated from amino acid sequences of selected land plant species (model of substitution: WAG+F+I+G4, lnL: -18520.157). Coloured labels indicate orthologs of *Calypogeia fissa* (blue), and the known orthologs of Marchantia paleacea and Medicago truncatula (green). Closed circles on branches indicate SH-like support values > 0.8. Scale bars are in expected substitutions per site. See Table S8 for species list and abbreviations.

Maximum-likelihood tree estimated from amino acid sequences of selected land plant species (model of substitution: WAG+F+I+G4, lnL: -18520.157). Coloured labels indicate orthologs of *Calypogeia fissa* (blue), and the known orthologs of *Marchantia paleacea* and *Medicago truncatula* (green). Closed circles on branches indicate SH-like support values > 0.8. Scale bars are in expected substitutions per site. See Supplementary Table S8 for species list and abbreviations.

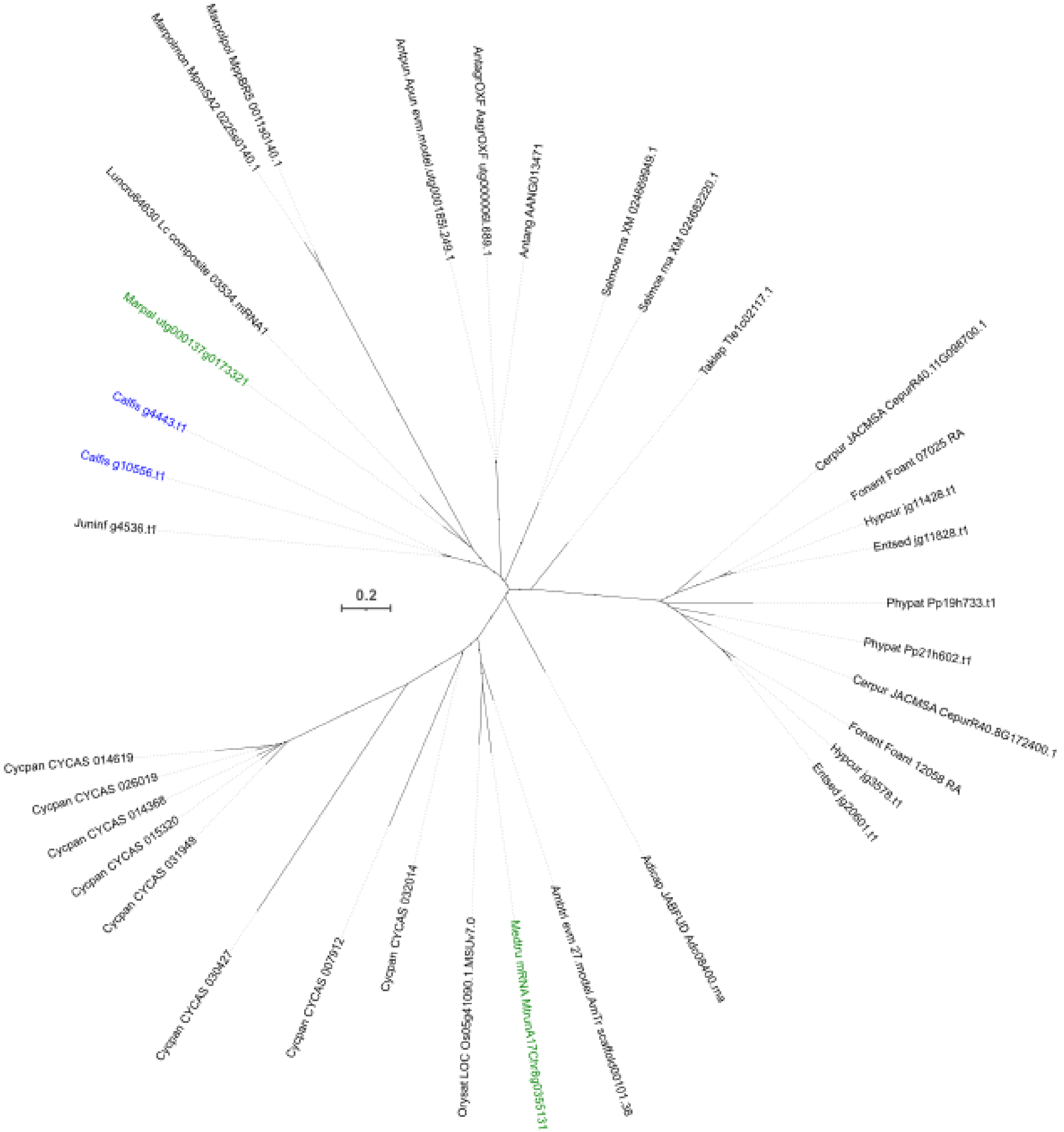


**Fig. S3. Phylogenetic tree of *CCaMK.*** Maximum-likelihood tree estimated from amino acid sequences of selected land plant species (model of substitution: JTT+G4, lnL: -13493.574). See legend of Figure S2 for description of the tree.

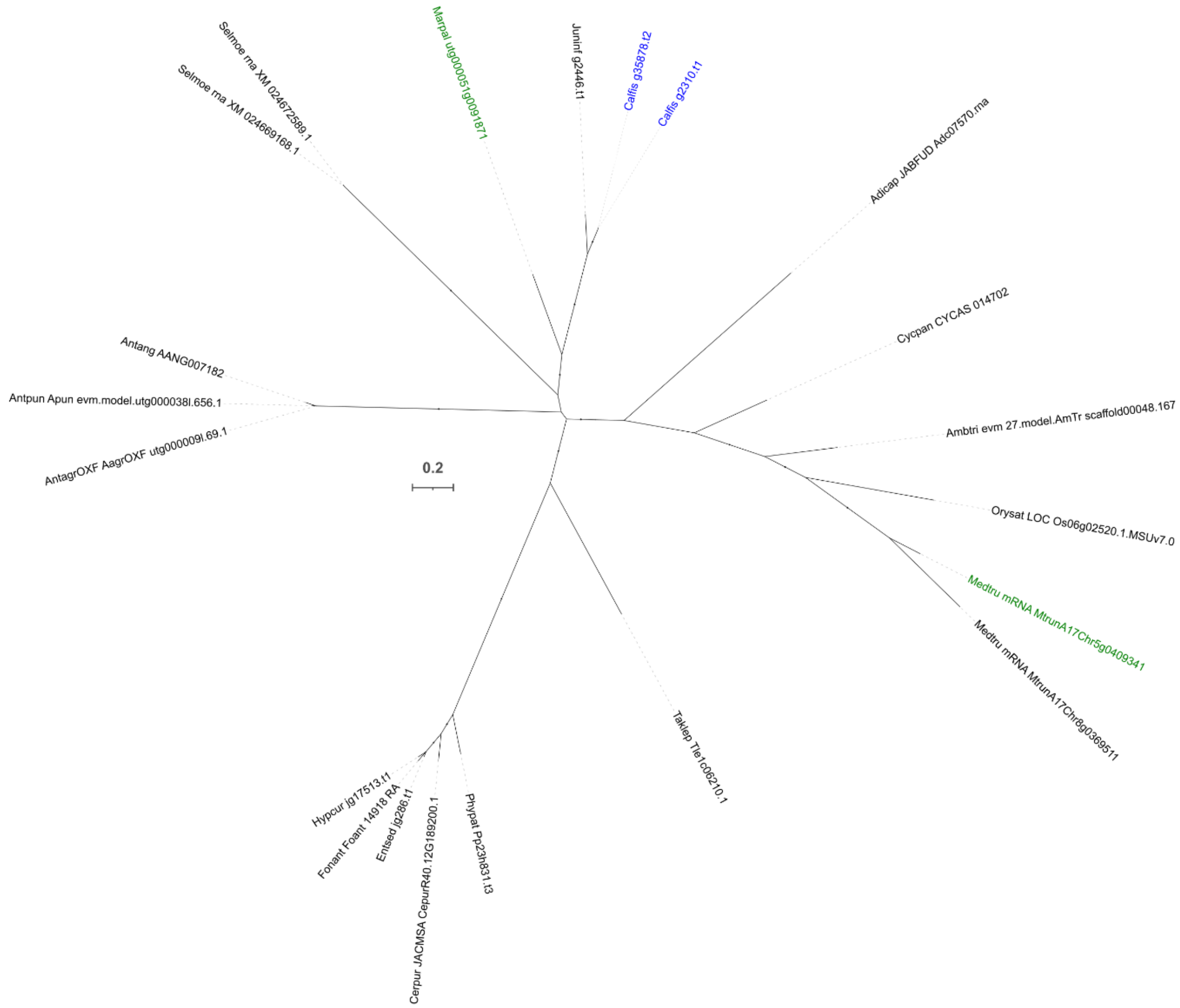


**Fig S4. Phylogenetic tree of *CYCLOPS*.** Maximum-likelihood tree estimated from amino acid sequences of selected land plant species (model of substitution: Q.mammal+F+I+G4, lnL: -16941.399). See legend of Figure S2 for description of the tree.

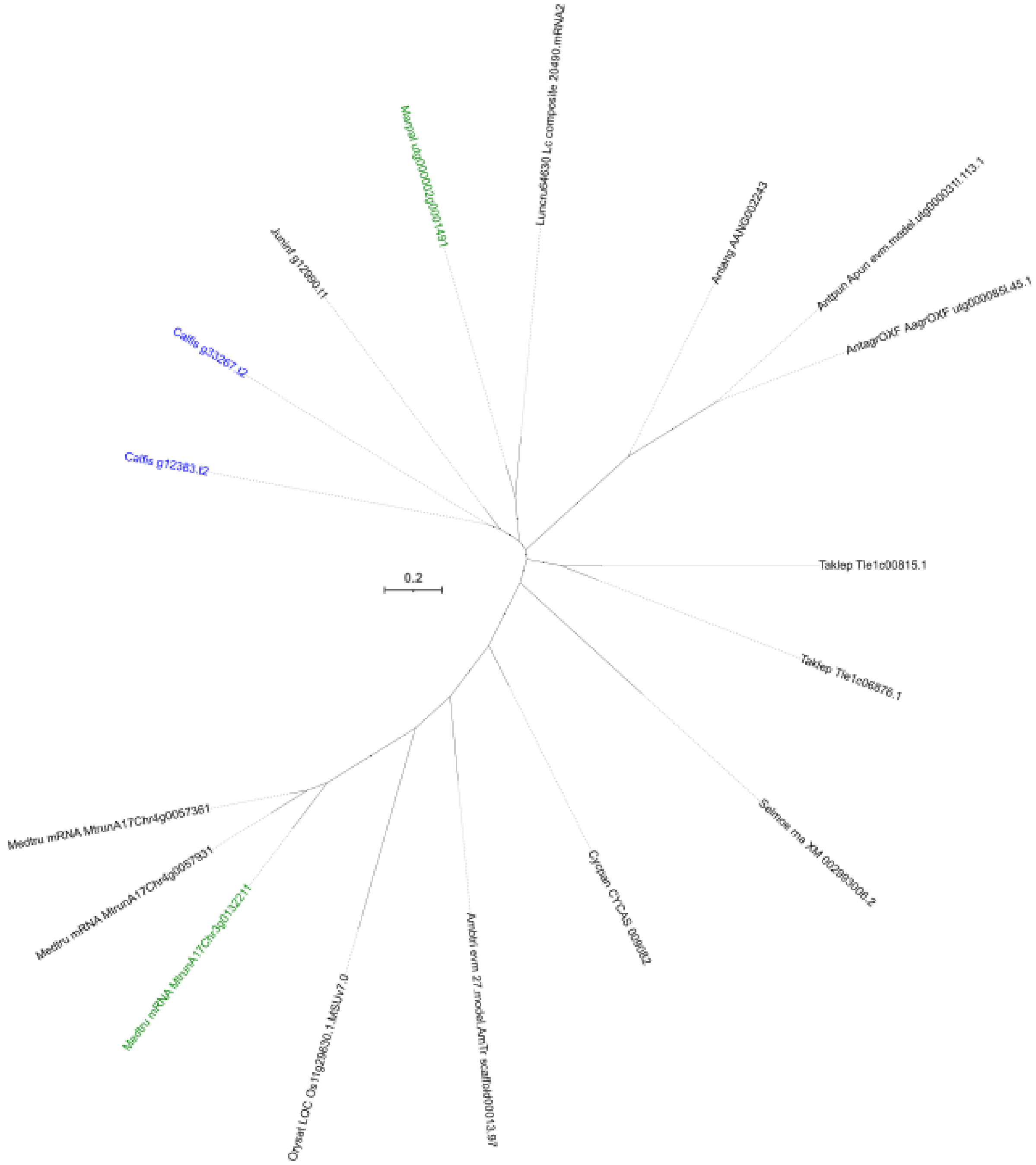


**Fig. S5. Phylogenetic tree of *EPP1*.** Maximum-likelihood tree estimated from amino acid sequences of selected land plant species (model of substitution: WAG+F+I+G4, lnL: -18520.157). See legend of Figure S2 for description of the tree.

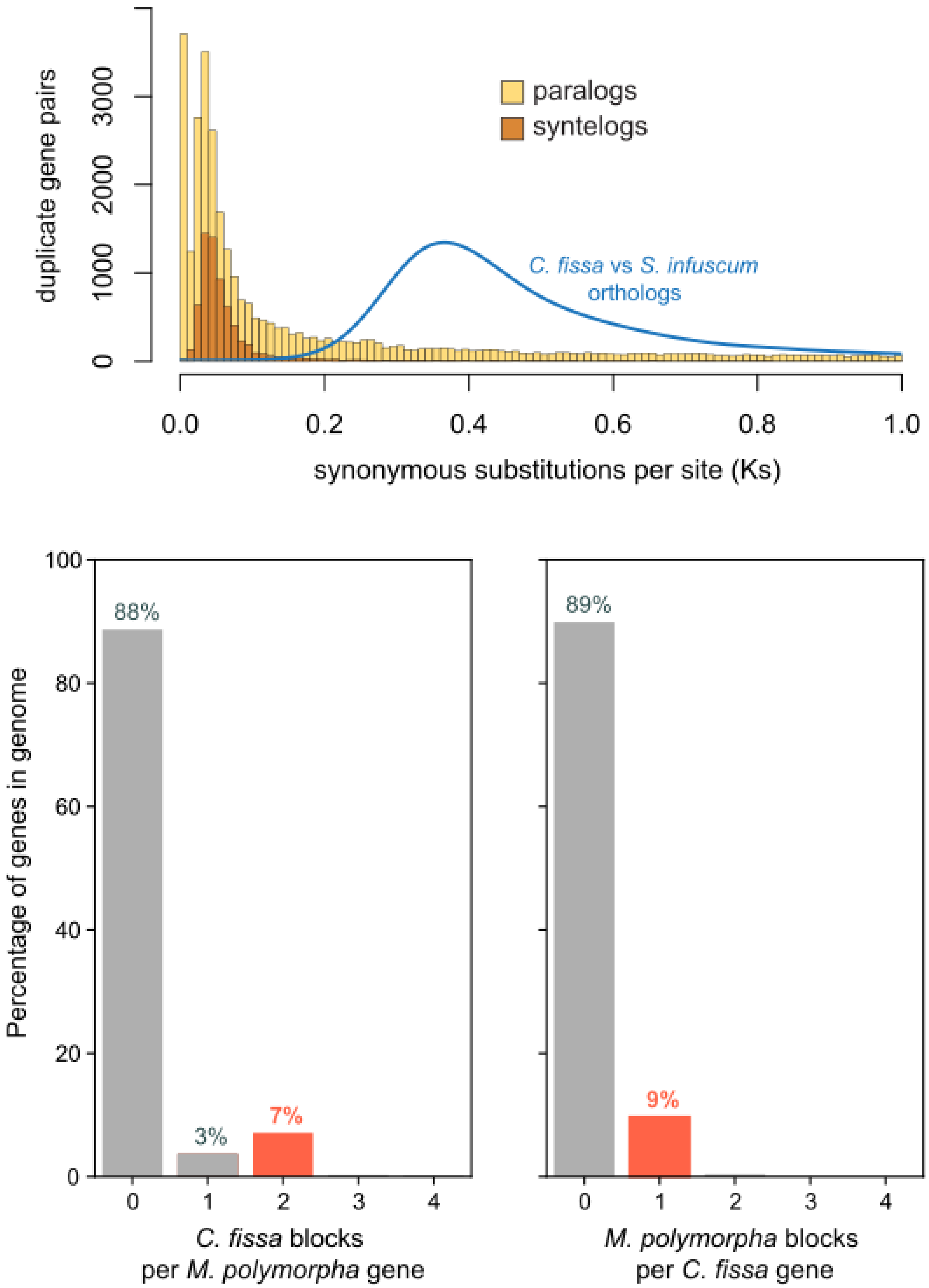


**Fig. S6.**
**Orthologous Ks distribution.** Orthologous Ks distribution in *Calypogeia fissa* and *Solenostoma infuscum* and reciprocal gene block compraison between *C. fissa* and *M. polymorpha* provides evidence for a recent Whole Genome Duplication (WGD) in *C. fissa*.

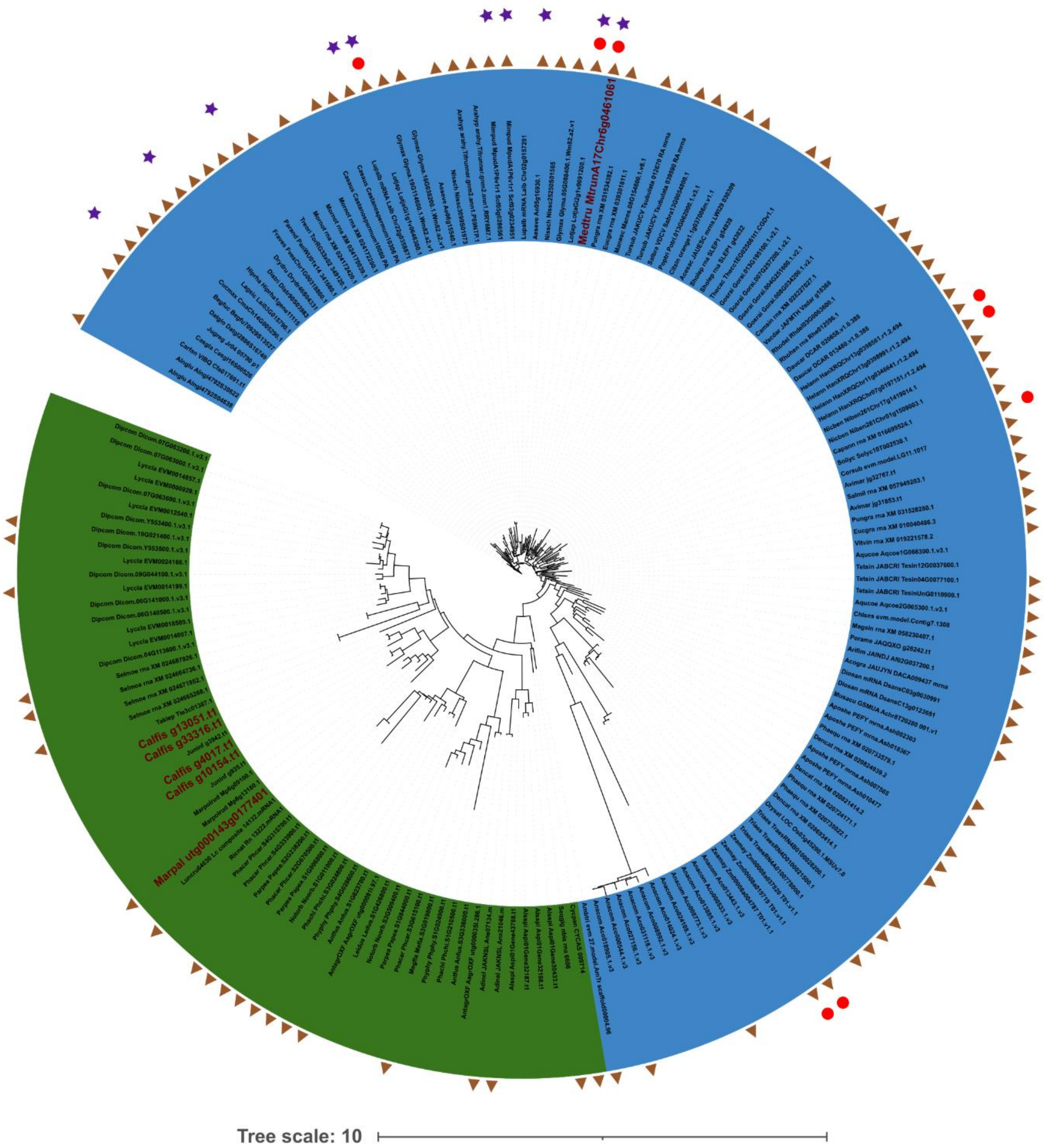


**Fig. S7. Phylogenetic tree of *VAPYRIN*.** Maximum-likelihood tree estimated from amino acid sequences of selected land plant species (model of substitution: Q.plant+I+G4, lnL: -71848.719). Angiosperms are in blue and bryophytes in green. *Medicago truncatula*, *Marchantia paleacea* and *Calypogeia fissa* genes are highlighted in red. Presence of the ‘GCCGGC’ motif (brown triangles), up-regulated genes during AMS (red circles), and up-regulated genes during RNS (purple stars) are shown.

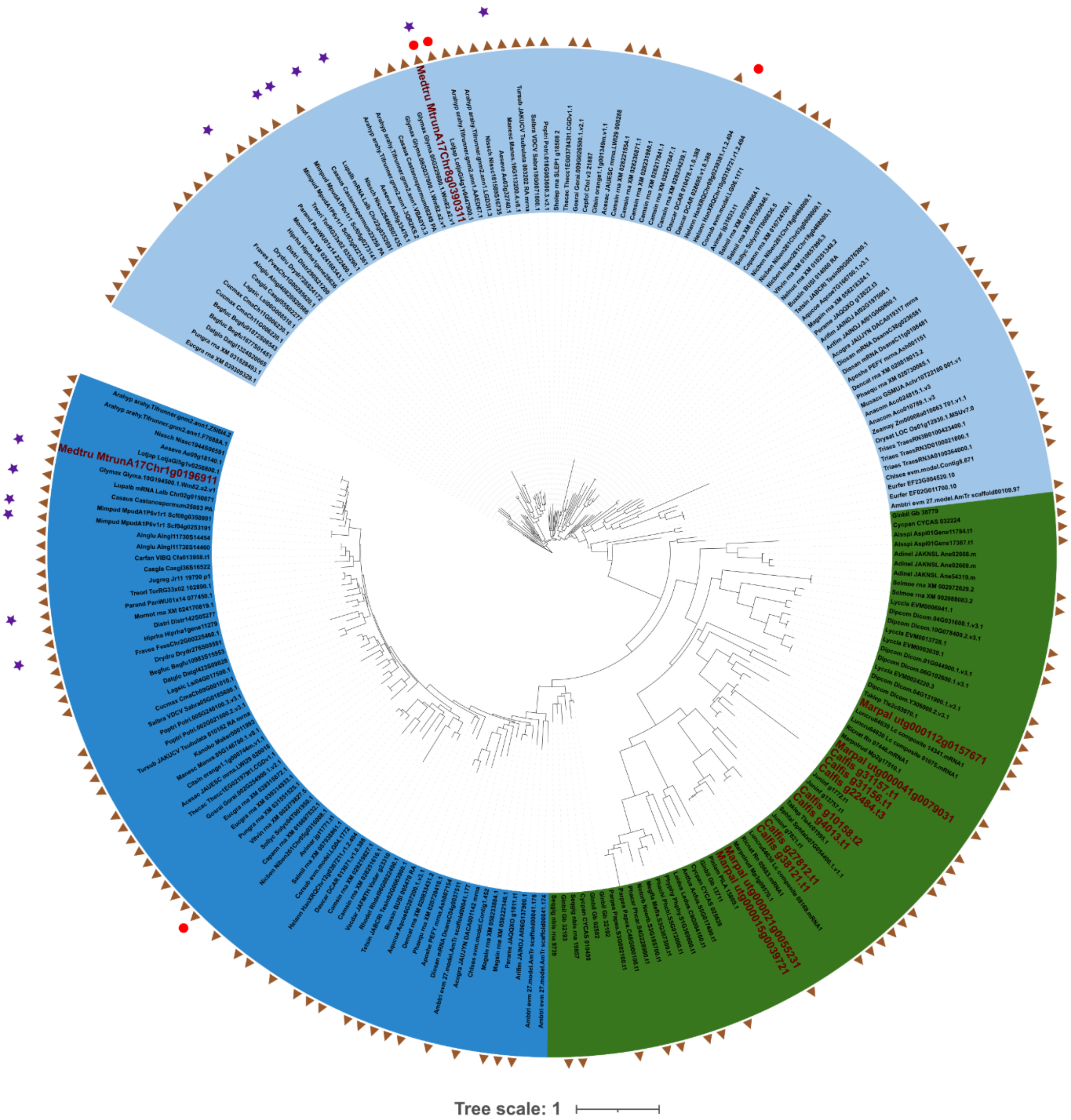


**Fig. S8. Phylogenetic tree of *LIN*.** Maximum-likelihood tree estimated from amino acid sequences of selected land plant species (model of substitution: Q.mammal+F+I+G4, lnL: -255261.449). Angiosperms are in blue and bryophytes in green. *Medicago truncatula*, *Marchantia paleacea* and *Calypogeia fissa* genes are highlighted in red. Presence of the ‘GCCGGC’ motif (brown triangles), up-regulated genes during AMS (red circles), and up-regulated genes during RNS (purple stars) are shown.

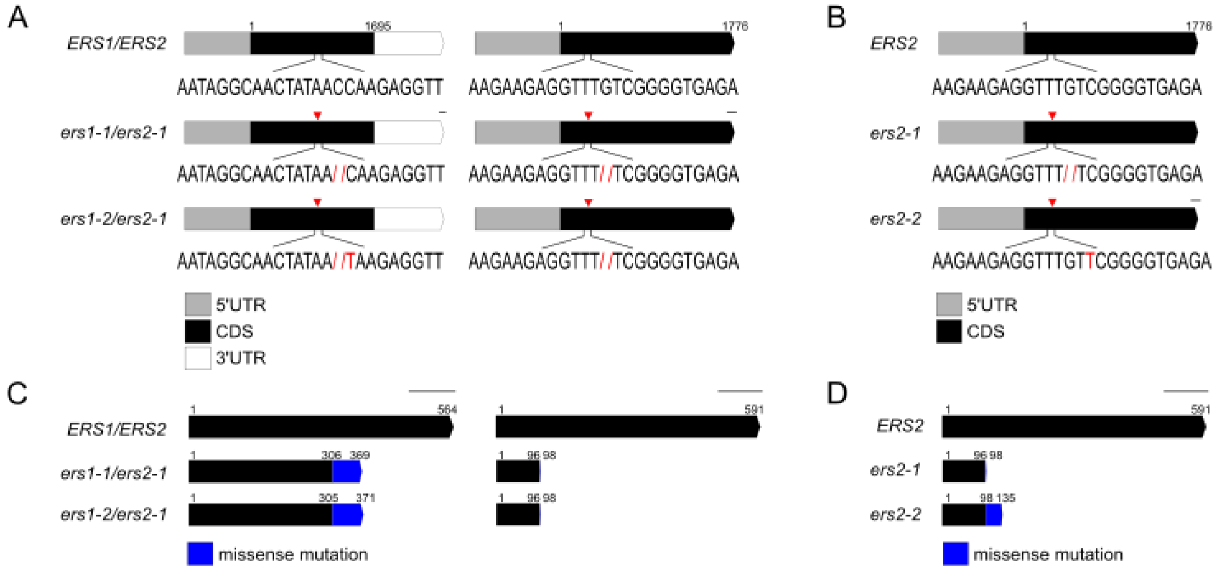


**Fig. S9. Gene models and predicted proteins of *Mpaers* mutants.** (A) Schematic overview of mRNA structure of wildtype Mpa*ERS* and mutated versions in Mpa*ers* double mutants. Red arrows indicate the site of mutations. (B) Schematic overview of mRNA structure of wildtype Mpa*ERS2* and mutated versions in Mpa*ers2* single mutants. Red arrows indicate the site of mutations. (C) Schematic overview of predicted proteins of wildtype Mpa*ERS* and mutated versions in Mpaers double mutants. (D) Schematic overview of predicted proteins of wildtype Mpa*ERS2* and mutated versions in Mpa*ers2* single mutants. Gene models were generated with http://wormweb.org/exonintron.

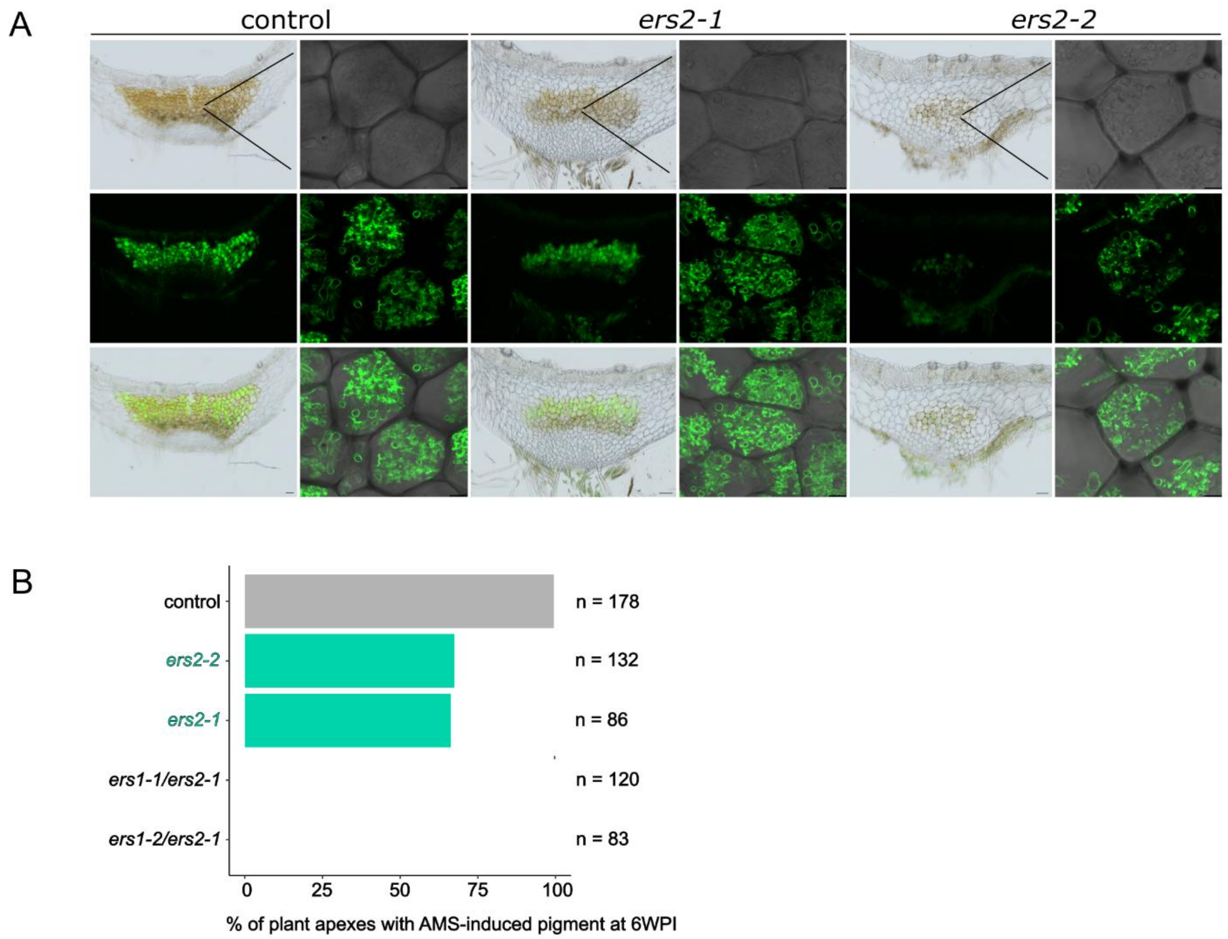


**Fig. S10. Phenotypic characterization of *Mpaers2* single mutants during AMS.** (A) Representative images of control plants transformed with an empty vector and Mpa*ers2* single mutants inoculated with *R. irregularis* at 6 weeks post inoculation. Green channel: WGA-Alexa 488 staining of fungal structures. Scale bar 100 µM or 10 µM. (B) Scoring of plant apexes with AMS-induced pigment at 6 weeks post inoculation with *R. irregularis* in control plants transformed with an empty vector, Mpa*ers* double and Mpa*ers2* single mutants. Combined results of 4 independent experiments are shown.

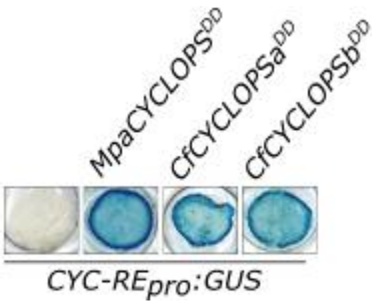


**Fig. S11. CYCLOPS transactivation assays in *Nicotiana benthamiana.*** *N. benthamiana* leaves expressing *CYC-REpro:GUS* alone or together with constructs mimicking CYCLOPS phosphorylation (*CYCLOPS*$^{DD}$) from *M. paleacea* (*MpaCYCLOPS*$^{DD}$) and *C. fissa* (*CfCYCLOPSa*$^{DD}$, *CfCYCLOPSb*$^{DD}$). GUS staining was performed at 48HPI and is shown in blue.

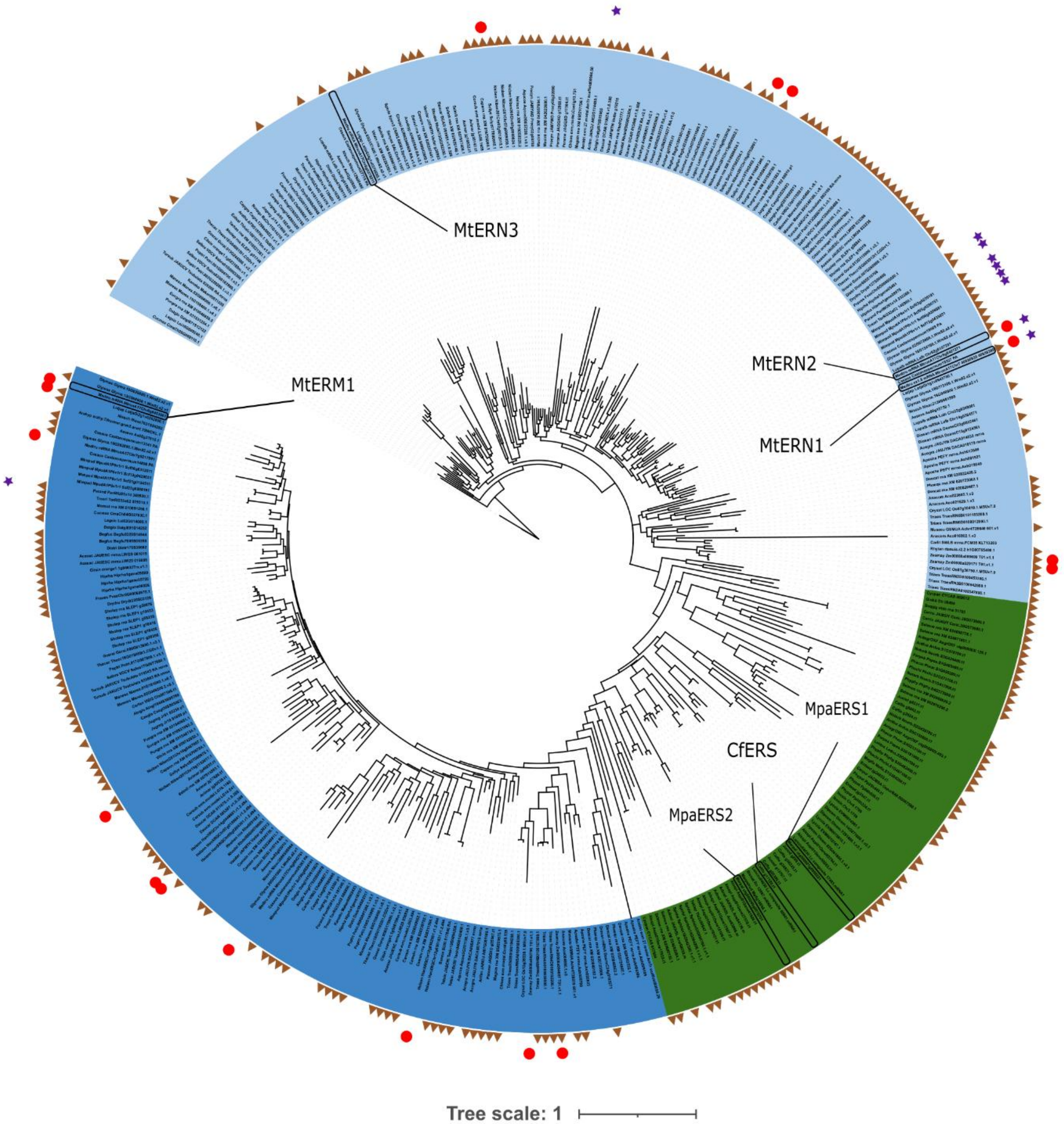


**Fig. S12. Phylogenetic tree of *ERS*.** Maximum-likelihood tree estimated from amino acid sequences of selected land plant species (model of substitution: JTT+I+G4, lnL: -140845.098). The 2 clades of angiosperms are in blue and bryophytes in green. *Medicago truncatula*, *Marchantia paleacea* and *Calypogeia fissa* ERF-like genes are pointed. Presence of the *CYC-RE* motif (brown triangles), up-regulated genes during AMS (red circles), and up-regulated genes during RNS (purple stars) are shown.

| | *C. fissa* | *S. infuscum* |
|---|---|---|
| **Genome size (bp)** | 763616257 | 383453254 |
| **Number of sequences** | 82 | 4616 |
| **N50** | 15851540 | 244190 |
| **L50** | 14 | 480 |
| **GC content (excluding Ns) (%)** | 43,3 | 43,1 |
| **Number of predicted CDS** | 38150 | 15468 |
| **Genome completion** | C:96.0%[S:2.1%,D:93.9%],F:0.5%,M:3.5% | C:95.8%[S:92.5%,D:3.3%],F:0.5%,M:3.8% |
| **Annotation completion** | C:96.5%[S:1.9%,D:94.6%],F:0.7%,M:2.8% | C:94.8%[S:91.1%,D:3.8%],F:0.7%,M:4.5% |

**Table S1. Genome assembly statistics for the leafy liverworts *Solenostoma infuscum* and *Calypogeia fissa.***

| | | Plow* Nstandard | Plow Nhigh | Phigh** Nstandard | Phigh Nhigh | Phigh Nlow | Plow Nlow | Pstandard Nhigh |
|---|---|---|---|---|---|---|---|---|
| | g $mol^{-1}$ | | | | g $L^{-1}$ | | | |
| $KNO_3$ | 101 | N/A | 7.5 | N/A | 7.5 | N/A | N/A | 7.5 |
| $Ca(NO_3)_2•4H_2O$ | 236 | 1.78 | 9.5 | 1.78 | 9.5 | 0.24 | 0.36 | 9.5 |
| $NaH_2PO_4•2H_2O$ | 156 | 0.02 | 0.02 | 4.68 | 4.68 | 3.9 | 0.02 | 2.34 |
| $MgSO_4•7H_2O$ | 246 | 5 | 5 | 5 | 5 | 5 | 5 | 5 |
| $K_2SO4$ | 174 | 6.46 | 6.46 | 6.46 | N/A | 6.46 | 6.46 | 6.46 |
| $CaCl_2•2H_2O$ | 147 | 2.4 | 2.4 | 2.4 | N/A | N/A | 2.84 | 2.4 |

These macronutrient solutions were used to prepare the working solution by combining them with micronutrient and iron-containing solutions. This mixture was then applied into microboxes for conducting the nutrient supply experiments. See the Methods section for further details.
*Permissive and **non-permissive nutrient conditions for ericoid mycorrhizae.

**Table S2. Chemical composition of macronutrient solutions used in this study.**

**Table S3. InterPro terms enrichment associated to up-regulated genes in permissive condition.**

**Table S4. InterPro terms enrichment associated to down-regulated genes in permissive condition.**

**Table S5. InterPro terms enrichment associated to up-regulated genes in non-permissive condition.**

**Table S6. InterPro terms enrichment associated to down-regulated genes in non-permissive conditions.**

**Table S7. Orthogroups containing up-regulated genes of *C. fissa* in permissive condition only and of *M. paleacea***

**Table S8. List of species used in the phylogenetic and promoter analyses**

**Table S9. Summary of up and down-regulated *ERS*, *VAPYRIN*, and *LIN* in response to arbuscular mycorrhiza or root nodule nitrogen-fixing symbioses in angiosperms.**

**Table S10. Summary of putative regulatory motifs identified in promoters of *ERS*, *VAPYRIN* and *LIN*.**

**Table S11. Plasmids and sgRNA sequences used in this study**

**Table S12. Source of data used to re-estimate genes up-regulated during AMS**
